# Environmental drivers of *Ixodes ricinus* tick population dynamics: mechanistic modelling using longitudinal field surveys and climate data

**DOI:** 10.1101/2024.06.06.597751

**Authors:** Younjung Kim, Benoît Jaulhac, Juan F Vesga, Laurence Zilliox, Nathalie Boulanger, W John Edmunds, Raphaëlle Métras

## Abstract

*Ixodes ricinus* is the primary vector for Lyme disease and tick-borne encephalitis across Europe. Despite playing a critical role in disease transmission dynamics, the environmental drivers of its complex life cycle have not been quantified using real-world data. To address this gap, we fitted a unique mechanistic model to a detailed 10-years longitudinal dataset from four sites in Northern France, where *I. ricinus* is abundant and Lyme disease and tick-borne encephalitis have been reported for decades, within a Bayesian framework. By incorporating key demographic processes and the influence of environmental conditions on these processes, our model estimated oviposition, hatching, and moulting rates across a range of temperature or saturation deficit, as well as questing and vertebrate host contact rates. Notably, moulting peaked at 14.2°C (95%HDI: 12.5–16.1°C), substantially lower than commonly suggested by laboratory-based studies, whereas oviposition and hatching peaked at 24.4°C (95%HDI: 10.9– 27.2°C) and 24.7°C (95%HDI: 17.8–27.2°C), respectively. Furthermore, vertebrate host contact rates significantly varied between the four study sites, with one site presenting up to 2.90 (95%HDI: 2.15–3.86) times higher contact rates than the other three sites. Additionally, we showed the importance of diapause in reproducing the observed seasonal population dynamics. For ticks overwintering through diapause, moulting in spring more accurately matched the predominantly unimodal questing activity patterns observed, compared to moulting in summer. Finally, model projections under several climate change scenarios indicated decreasing tick abundance trends over the next two decades. This study provides a foundation for models of *I. ricinus*-borne pathogen transmission and can be adapted to other *Ixodidae* populations of public health significance.

## Introduction

*Ixodes ricinus* is the most widespread tick species across Europe, transmitting pathogens that pose significant risks to both public and animal health (1). These pathogens include *Borrelia burgdorferi* sensu lato causing Lyme disease—the most prevalent vector-borne disease in Europe (2)—and tick-borne encephalitis virus (TBEV), which can lead to fatal neurological disorders with a growing incidence throughout the continent (3).

*Ixodidae* ticks including *I. ricinus* undergo a complex life cycle comprised of a series of key demographic processes, including egg-laying (oviposition), larvae emerging from eggs (hatching), larvae, nymphs and adults seeking vertebrate hosts (questing), feeding on vertebrate hosts (host contact), and progressing to the next life stage (larva-to-nymph and nymph-to-adult moulting). These processes may occur with or without delays in development (developmental diapause) or questing (behavioural diapause) depending on environmental conditions (4–7).

Understanding the mechanisms driving these processes is crucial for unravelling tick population dynamics. This, in turn, enables the assessment of current and future tick-borne disease incidence in animal hosts and humans, as well as the development of effective public and animal health strategies.

Mechanistic modelling provides an ideal framework for understanding tick population dynamics by enabling explicit descriptions of tick progression through key demographic processes. For *Ixodes* ticks, although few in number, mechanistic models have been developed to capture the seasonality of questing tick density by accounting for the impact of temperature on tick demographic processes (8–11), and have been further expanded to explore the transmission dynamics of tick-borne pathogens (12–14). While providing valuable insights into the population dynamics of *Ixodes* ticks, the application of these models has been limited for several reasons. Firstly, most simulation studies have relied on parameter values derived from controlled laboratory or field settings (5, 15–17). However, significant spatiotemporal variations in climate and *Ixodes* ticks’ adaptability to a wide range of climatic conditions (18, 19) suggest that those parameter values do not fully represent how ticks have adapted to their dynamic natural environments (20). Secondly, relying on simulations with fixed parameter values fails to capture the variability inherent in observed data (21). This restricts the value of mechanistic models since understanding the extent of data variability can not only provide insights into the unobserved ecological processes but also allow more reliable predictions of tick population sizes and the burden of tick-borne pathogens.

Here, we collected longitudinal data on questing *I. ricinus* nymph and adult density from 2013 to 2022 in four distinct sites (Dannemarie, Illkirch, Murbach, and Niedermunster) in the Alsace region of Northeast France, where Lyme disease and tick-borne encephalitis have been reported for decades (22–24). We developed and fitted a mechanistic model to this longitudinal data using a Hamiltonian Monte Carlo (HMC) Bayesian algorithm, to estimate key demographic parameters related to tick development, questing, and vertebrate host contact activities, and to test hypotheses regarding the following two key features in the life cycle of *I. ricinus* (Figs. 1 and 2). Firstly, we assessed the functional relationship between climatic conditions and tick development by estimating the daily rates of oviposition, hatching, larva-to-nymph moulting, and nymph-to-adult moulting according to temperature or saturation deficit, key environmental factors known to influence tick development (4) (Theme 1 hypotheses). Secondly, we investigated the impact of developmental diapause—a delay between tick feeding and moulting—on the emergence of newly moulted, and thus questing ticks (Theme 2 hypotheses). This was done by evaluating model fits under different lengths of developmental diapause based on day length, which has been suggested to modulate diapause (4). Finally, based on the best-fitting model, we assessed the potential impact of climate changes for the next 20 years on *I. ricinus* abundance in the study region by using climate change scenarios represented by Representative Concentration Pathways (RCP) (25).

**Figure 1.**
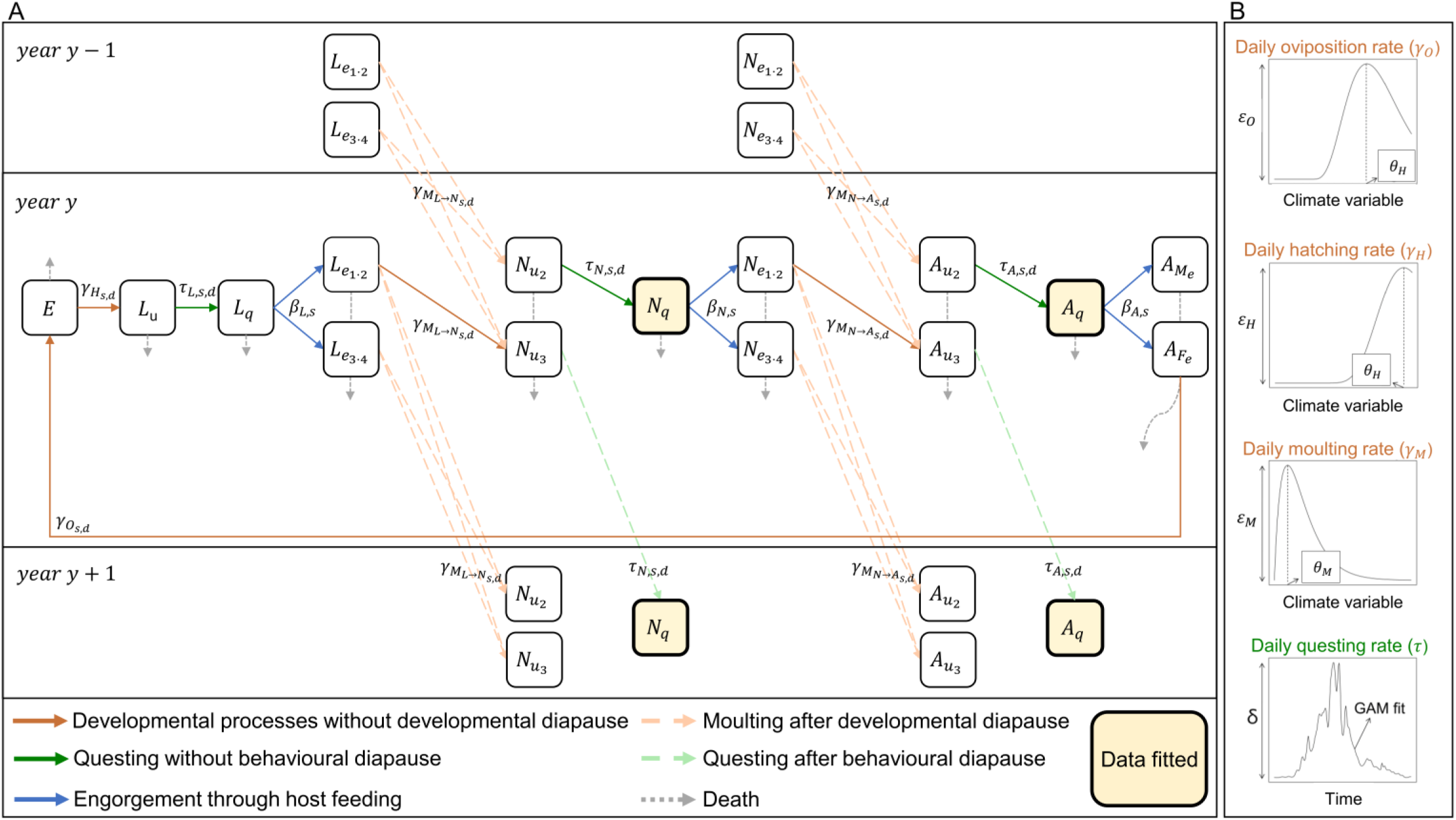
Schematic description of Ixodes ricinus population dynamics model. See Tables 1 and 2 for the definition of parameters shown in this figure. (A) Transition of ticks through life stages. Square boxes represent the compartments of ticks in various life stages: E=eggs, L=larvae, N=nymphs, A=adult, A_M_=male adults, and A_F_=female adults. Subscripts u, q, and e represent unfed, questing, and engorged statuses, respectively. Square boxes with bold outline and ivory colour represent questing nymph and adult populations to which the model was fitted. Arrows between the compartments indicate the daily rates of developmental (brown), questing (green), or host contact (blue) processes, with the subscripts s, d indicating study site and date, respectively. Dotted pale brown arrows represent a delay in moulting through developmental diapause, and dotted pale green arrows represent a delay in questing through behavioural diapause. Grey dotted arrows represent mortality through natural death. (B) Parameters estimated for daily oviposition, hatching, moulting and questing rates. For daily oviposition, hatching, and moulting rates, the climatic condition (temperature or saturation deficit) at their peaks (𝜃_𝑂_, 𝜃_𝐻_, and 𝜃_𝑀_) and the corresponding maximum daily rates (𝜀_𝑂_, 𝜀_𝐻_, and 𝜀_𝑀_) were estimated assuming that these rates followed the shape of a Gamma distribution under Theme 1 alternative hypothesis. Under Theme 1 null hypothesis, these developmental rates were assumed to follow the inverse of the power-law functions estimated by Ogden, Lindsay, Beauchamp, Charron, Maarouf, O’Callaghan, Waltner-Toews and Barker (*2*). In both baseline and alternative hypotheses, daily questing rates were obtained by scaling temporal trends in questing tick density, estimated by the best-fitting generalised additive model (GAM), by parameter 𝛿.

**Figure 2.**
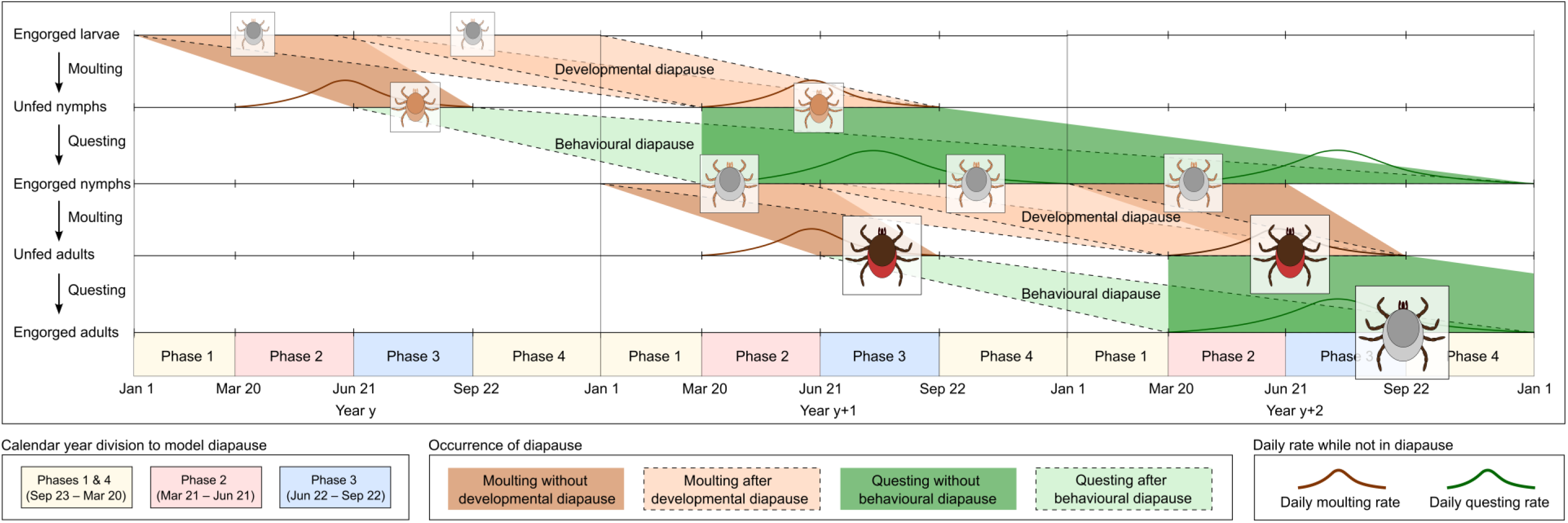
Occurrence of developmental and behavioural diapauses in relation to the timing of moulting and questing. Upper four rows illustrate the transition of ticks from engorged larvae to engorged adults, through moulting and questing, with or without experiencing a developmental or behavioural diapause. Bottom row displays the division of each calendar year into four phases, which govern tick demographic processes. Under Theme 2 null hypothesis, ticks that had their blood meal during Phase 2 or earlier began moulting from Phase 3 of the same year (dark brown shades). Some ticks that did not moult in Phase 3 began moulting in Phase 2 of the following year after a developmental diapause (pale brown shades with a dashed border). Additionally, ticks that had their blood meal in Phase 3 began moulting from Phase 2 of the following year after a developmental diapause (pale brown shades with a dashed border). Under Theme 2 alternative hypothesis, these ticks began moulting from Phase 3 of the following year instead (not shown in the figure). Regarding questing, ticks that moulted after a developmental diapause began questing from Phase 2 of the year they moulted (dark green shades), whereas ticks that moulted in Phase 3 without a developmental diapause began questing from Phase 2 of the following year after a behavioural diapause (pale green shades with a dashed border). While moulting and questing were permitted under these conditions, they were subject to occur according to daily moulting (brown lines) or questing (green lines) rates, as estimated under Theme 1 hypotheses.

**Table 1.**
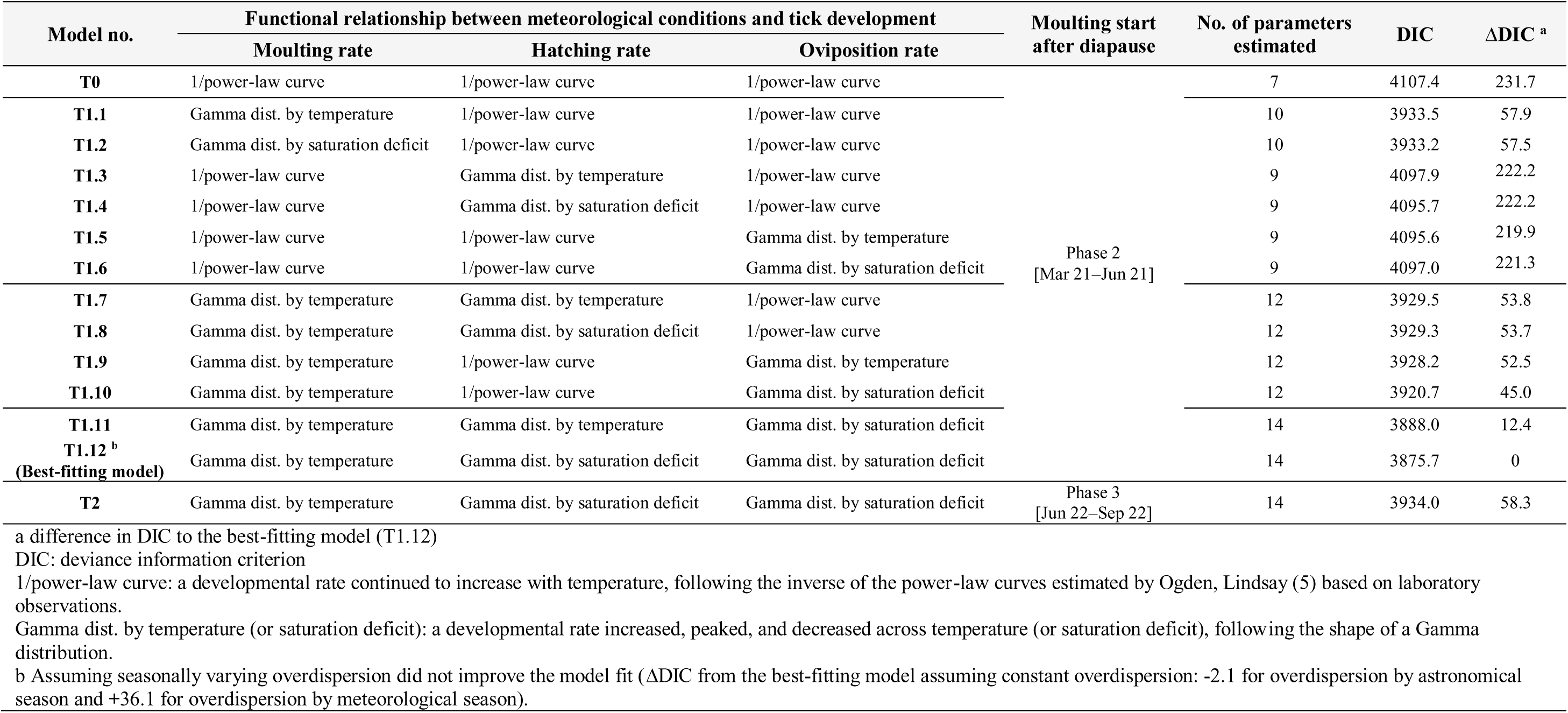
Comparison of mechanistic models under different sets of Theme 1 and 2 hypotheses.

**Table 2.** Parameters estimated by the mechanistic model. Notations, definitions, priors, and assumptions.

| Notation | Definition | Prior | Assumption |
| --- | --- | --- | --- |
| **Daily questing rate** |  |  |  |
| $\tau_{l,s,d}$ | Daily questing rate by life stage, site, and day | Computed from estimated parameter $\delta$ and GAM fit values | The daily questing rate followed the trends in GAM fit values, accounting for climatic variables (see Appendix S1). Larvae and nymphs were assumed to have same questing rates. |
| $\delta$ | A scaling parameter for the daily questing rate | Uniform (0, 0.143) | The upper prior limit was set to ensure that ticks remain unfed for at least 7 days after moulting, to account for the hardening process (4). |
| **Daily host contact rate** |  |  |  |
| $\beta_{l,s}$ | Daily host contact rate by life stage and site | Computed from estimated parameter $\rho$ and assumed baseline host contact rate $\phi$, as per Eq. 2 | For each life stage and study side, $\beta$ was derived by scaling the baseline daily host contact rate (Table S1) using the parameter $\rho$. |
| $\rho_s$ | A scaling parameter for the daily host contact rates by site | Uniform (0, 11.1) | The upper prior limit is set to ensure that $\beta$ remains below 1. |
| **Developmental rate under Theme 2 alternative hypothesis** |  |  |  |
| $\gamma_{O_x}$<br>$\gamma_{H_x}$<br>$\gamma_{M_L \rightarrow N_x}$<br>$\gamma_{M_N \rightarrow A_x}$ | Daily oviposition, hatching, and larva-to-nymph and nymph-to-adult moulting rates at temperature or saturation deficit $x$ | Computed from estimated parameters $\theta$ and $\epsilon$, as per Eq. 1 | The daily developmental rate was assumed to vary with climatic conditions following the shape of a Gamma distributions. |
| $\theta_O$<br>$\theta_H$<br>$\theta_M$ | A temperature (or saturation deficit) at which a given daily developmental rate peaks following a gamma distribution shape | Temperature: Uniform (0, 27.3°C)<br>Saturation deficit: Uniform (0, 14.6) | The daily developmental rate was assumed to peak between 0°C (or a saturation deficit of 0) and the maximum temperature (or saturation deficit) observed during the study period. Larva-to-nymph and nymph-to-adult moulting rates were assumed to peak at the same temperature (or saturation deficit). |
| $\epsilon_O$<br>$\epsilon_H$<br>$\epsilon_{M_L \rightarrow N}$<br>$\epsilon_{M_N \rightarrow A}$ | A scaling parameter for daily oviposition, hatching, and larva-to-nymph and nymph-to-adult moulting rates | Uniform (0, 0.105)<br>Uniform (0, 0.066)<br>Uniform (0, 0.043)<br>Uniform (0, 0.057) | The peak daily developmental rates, estimated to occur within the observed temperature ranges according to the power-law functions of Ogden, Lindsay (5), were increased by an additional 25% to accommodate possibly species differences. These values were set as the upper prior limits. |
| **Parameters for model fitting based on log-likelihood values** |  |  |  |
| $\phi_{N_q}$ | Overdispersion for nymph in a negative binomial distribution | Uniform (0, 10) | The observed tick density was assumed to exhibit overdispersion, attributed to unknown heterogeneity in the data, such as unaccounted demography processes and sampling bias. |
| $\phi_{A_q}$ | Overdispersion for adults in a negative binomial distribution | Uniform (0, 10) | |

## Results

### Longitudinal *Ixodes ricinus* questing density data

In the four study sites, we collected questing *I. ricinus* nymphs and adults from 2013 to 2022. Within each study year, we conducted multiple sampling campaigns, primarily between March and November, resulting in a total of 368 sampling campaigns across the study sites (Fig. S1A). Each study site was sampled a median of 9 times a year (range: 7–11), totalling between 89 and 95 sampling campaigns during the study period. Across the study sites, questing *I. ricinus* nymphs and adults exhibited distinct annual peaks in the spring or early summer, with magnitudes that varied over the study years (Figs. 3 and 4). The highest questing tick density recorded was 426.25 per 100m^2^ for the nymphs (June 24, 2013, at the Murbach site) and 18 per 100m^2^ for the adults (May 18, 2016, at the Niedermunster site) (Figs. 3 and 4). Annual peak densities were predominantly observed between May 22 and June 17 for the nymphs and between May 18 and 30 for the adults (Figs. 3, 4, and S1B). This pattern demonstrated a primarily unimodal questing activity, consistent with previous observations in this region (19, 26).

**Figure 3.**
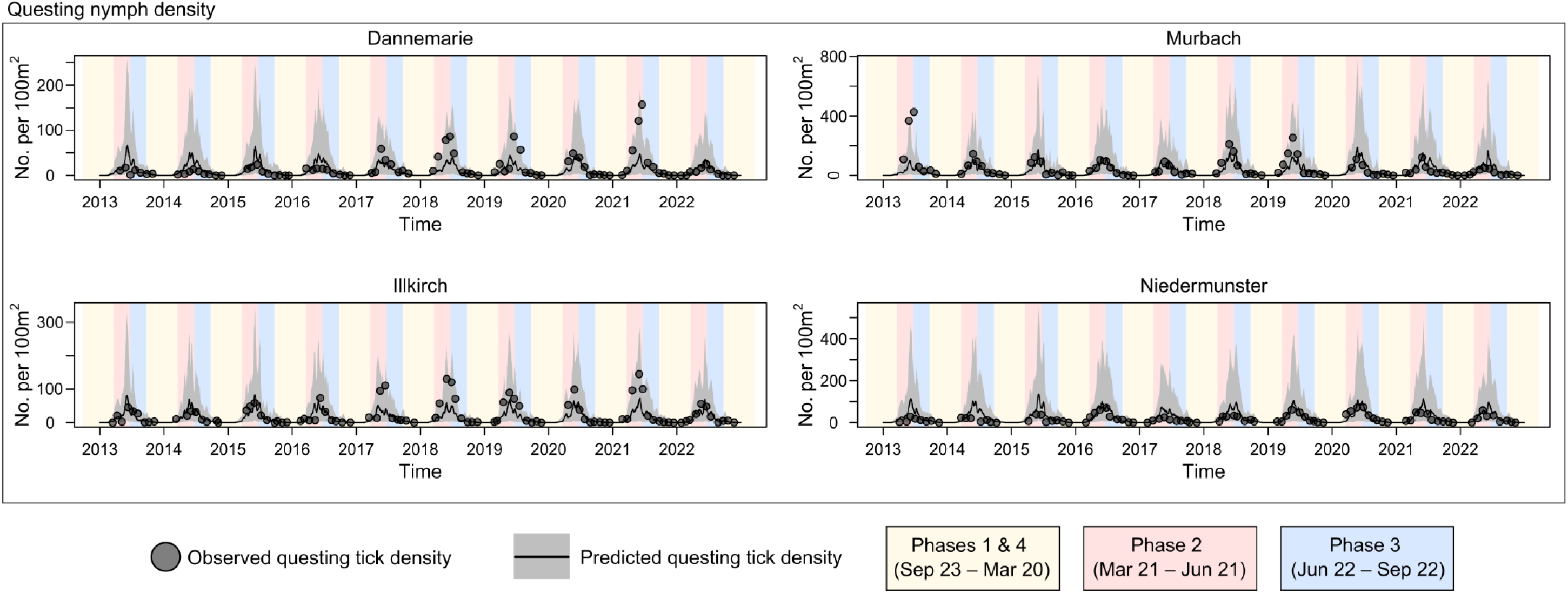
Observed and predicted longitudinal questing nymph density. Points represent the observed numbers of questing nymphs per 100m^2^, respectively. Black lines and grey shades represent the median and 95% percentile intervals of these metrics predicted by the best-fitting model.

**Figure 4.**
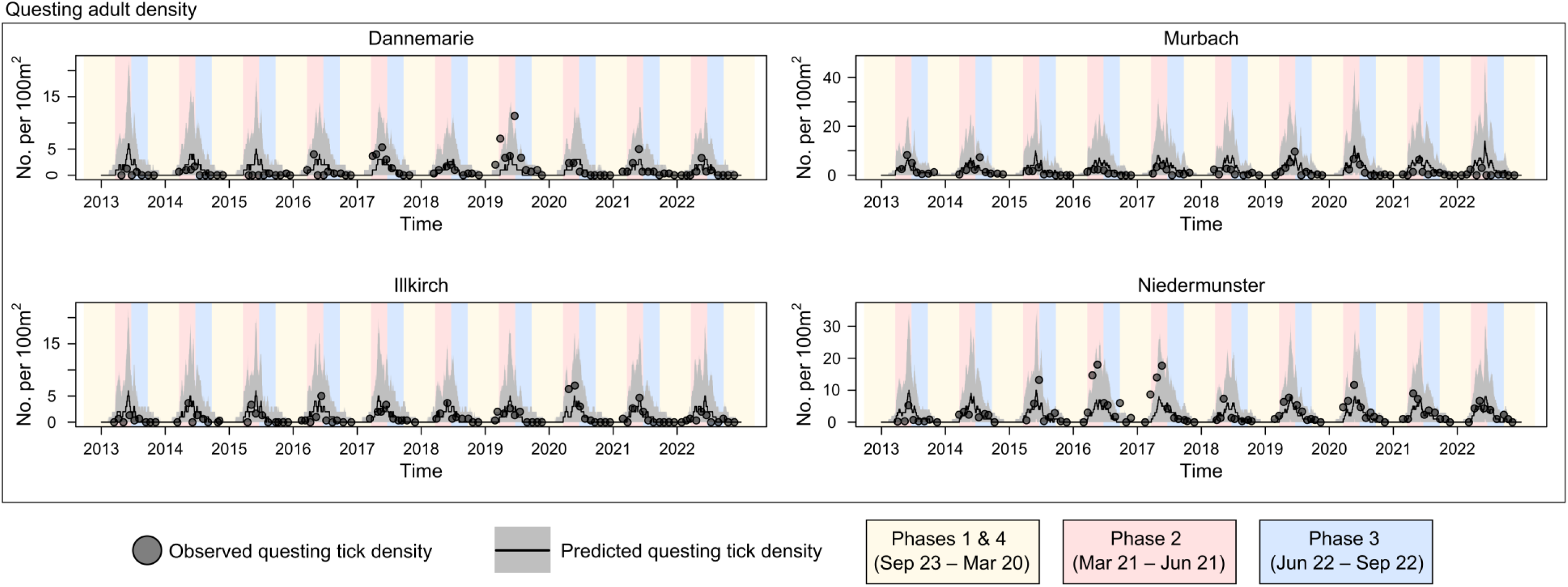
Observed and predicted longitudinal questing adult density. Points represent the observed numbers of questing adults per 100m^2^, respectively. Black lines and grey shades represent the median and 95% percentile intervals of these metrics predicted by the best-fitting model.

### Demographic model and tick density predictions

The progression of *I. ricinus* was modelled through four life stages: eggs (E), larvae (L), nymphs (N), and adults (A) (Fig. 1). Larvae, nymphs, and adults were further categorised into three stages: unfed (L_u_, N_u_, A_u_), questing (L_q_, N_q_, A_q_), and engorged (L_e_, N_e_, A_e_) compartments. Each transition between these three life stages required a blood meal from a vertebrate host and was influenced by environmental (climatic conditions) and ecological (developmental and behavioural diapauses) factors. In our model, these demographic processes were determined by the daily rates of developmental processes (i.e. oviposition, hatching, larva-to-nymph moulting, and nymph-to-adult moulting), questing (unfed ticks looking for a host), vertebrate host contact (questing ticks successfully feeding on a vertebrate host), and death, with or without developmental and behavioural diapauses (Fig. 2).

The model structure and underlying parameters are detailed in the Materials and Methods and Supporting Information. Briefly, each daily developmental rate was assumed to have a functional relationship with climatic conditions, either temperature or saturation deficit values. The model estimated parameters related to the underlying Gamma distribution and a scaling parameter that define this relationship. For the daily questing rates, we assumed that, for each life stage, they followed trends in questing tick density predicted by a generalised additive model, scaled by an estimated parameter. The daily baseline host contact rates were based on estimates from Dobson, Finnie and Randolph (8), with a separate scaling parameter estimated for each of the four study sites. Daily mortality rates for eggs, unfed larvae, unfed nymphs, and unfed adults were extracted from the literature (8, 27). Using these as baseline mortality rates, we made the following assumptions to maintain equilibrium in tick populations: (i) daily mortality rates increased linearly with population size above a specified threshold, accounting for theoretical survival probabilities between life stages (28) and (ii) once ticks had fed, they either progressed to the next life stage, laid eggs, or died within one year, reflecting the general pattern that fed ticks attempt to develop or oviposit when favourable weather conditions occur (4).

In the best-fitting model (T1.12 in Table 1), larva-to-nymph moulting and nymph-to-adult moulting were assumed to be influenced by temperature, while oviposition and hatching by saturation deficit, following the shape of a Gamma distribution (Table 1, see ‘Mechanistic model for tick population dynamics’ in the Materials and Methods for how these rates were modelled). This model explained the data significantly better than those assuming values from controlled laboratory settings for developmental rates (the inverse of power-law curves in Table 1). In simulation-based model assessments, our model effectively recovered parameter values used to generate synthetic tick density data, showing 95% HDI coverage ranging from 87.1% to 98.8% and no systematic bias patterns across the estimated parameters (Fig. S2) (see ‘Simulation-based model assessments and results’ in Appendix S1 Methods and Results for details).

The best-fitting model assumed that tick densities follow a negative binomial distribution with constant overdispersion over time. Assuming seasonally varying overdispersion did not explain the data significantly better than the best-fitting model (Table 1). Our model showed that in most sampling sessions across the sites, the median predicted densities aligned with the observed densities in most study years, with the 95th percentile intervals of the predicted densities encompassing the observed densities (96.7% for questing nymphs and 97.6% for questing adults) (Figs. 3, S3, and S4). On some occasions, the median predicted values did not reach the observed peak densities. Such excess variability was however captured by the overdispersion parameter (Figs. 3 and 4), estimated at 1.54 (95%HDI: 1.30–1.82) for questing nymphs and 2.13 (95%HDI: 1.38–2.97) for questing adults (Fig. S5F).

### Estimates of key demographic processes

The estimated peak of daily oviposition and hatching rates were 0.05 (95% highest density interval [HDI]: 0.01–0.10) and 0.05 (95%HDI: 0.02–0.07), occurring at daily mean saturation deficit values of 7.8 (95%HDI: 3.2–10.9) and 6.5 (95%HDI: 2.8–9.0), respectively (Figs. 5A–B and S5A–B). These saturation deficit values, being near the upper end of the values recorded during the study period (median: 2.2, 95th percentile interval [PI]: 0.5–8.5), indicated moderate to high drying power of the air. These rates exhibited similar variations across the range of daily mean temperature, with oviposition and hatching rates peaking at 24.4°C (95%HDI: 10.9–27.2°C) and 24.7°C (95%HDI: 17.8– 27.2°C), respectively, near the highest temperature recorded.

**Figure 5.**
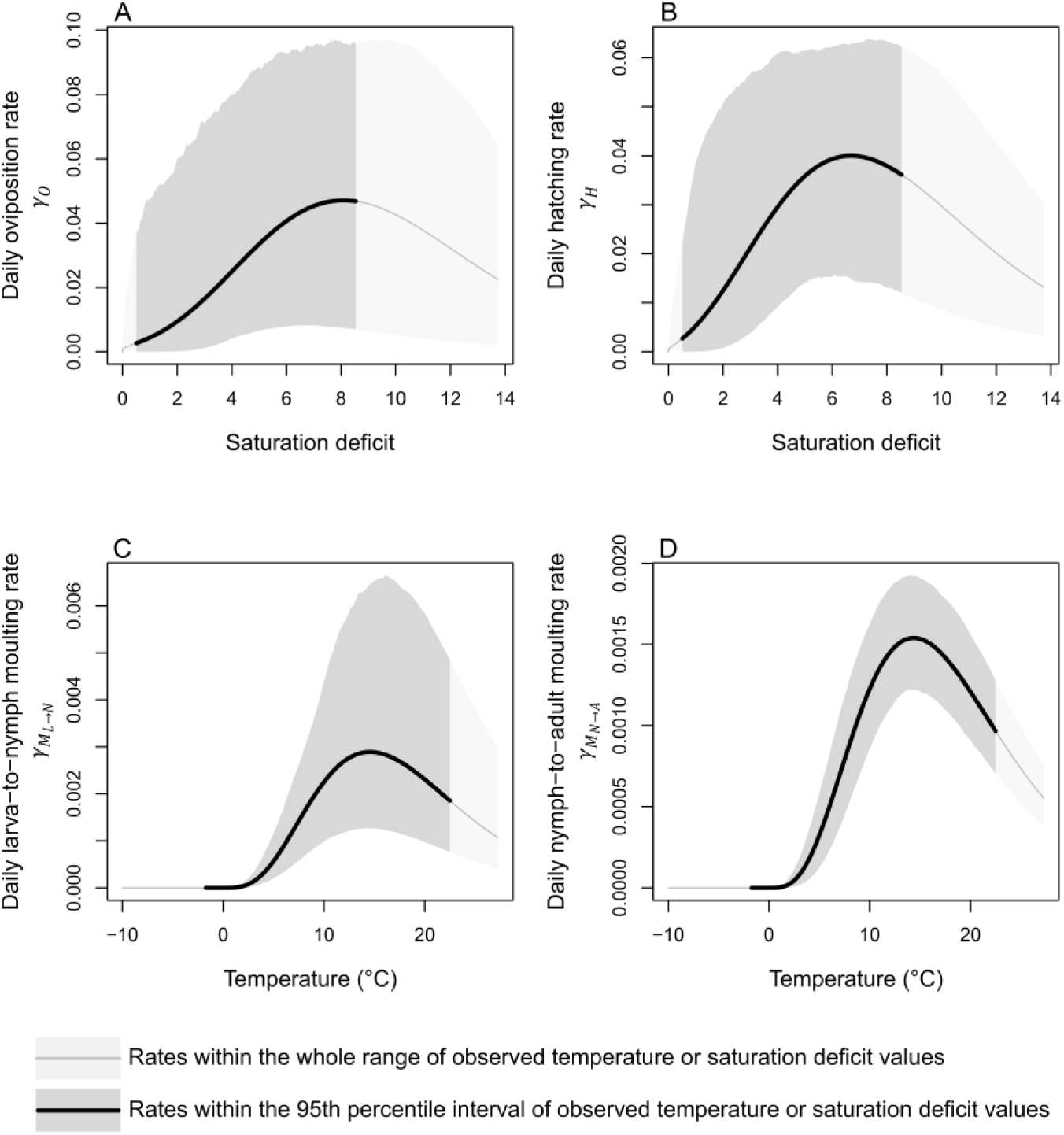
Functional relationship between climatic conditions and tick development. Daily developmental rates were estimated as a function of daily mean saturation deficit or temperature, under Theme 1 alternative hypothesis derived by the best-fitting model. It shows the functional relationship of daily (A) larva-to-nymph and (B) nymph-to-adult moulting rates with temperature, and the functional relationship of daily (C) oviposition and (D) hatching rates with saturation deficit. Lines and shades represent the median and 95th percentile interval of the developmental rates, respectively. Thick lines and dark shades represent the developmental rates within the 95^th^ percentile interval of the daily mean temperatures or saturation deficit values observed across the four sampling sites, while the thin lines and pale shades represent those rates within the whole range of the climatic variables observed.

Furthermore, we estimated that the highest larva-to-nymph and nymph-to-adult moulting daily rates were 3x10^-3^ (95%HDI: 1x10^-3^–5x10^-3^) and 2x10^-3^ (95%HDI: 1x10^-3^–2x10^-3^) respectively (Fig. S5B), both of which were significantly lower than the upper prior limits observed from laboratory observations (5). These peak moulting rates occurred at 14.2°C (95%HDI: 12.5– 16.1°C) (Figs. 5C–D and S5A), which was almost half the highest daily mean temperature recorded during the study period of 27.3°C.

The peak daily questing rate was 0.14 (median, 95%HDI: 0.12–0.14) (Fig. S5C), close to the upper prior limits established to allow time for ticks to harden their cuticle after moulting or hatching; at least 7 days of hardening were required for unfed ticks to begin questing. The median values of host contact rates ranged 0.12–0.35 for larvae, 0.18–0.51 for nymphs, and 0.23–0.64 for adults, across the study sites (Fig. S5E). We estimated the highest vertebrate host contact rates to be at the Murbach site, being 2.90 (95%HDI: 2.15–3.86), 2.64 (95%HDI: 1.91– 3.47), and 1.34 (95%HDI: 0.94–1.78) times higher than those at the Niedermunster, Dannemarie, and Illkirch sites, respectively (Fig. S5D).

Sensitivity analyses of the assumed daily baseline mortality and host contact rates were conducted. These analyses showed that changes in the assumed daily mortality rates (Figs. S6–S9) and baseline daily host contact rates (Figs. S10–S12) resulted in similar functional relationships between the developmental rates and climatic conditions, although the magnitude of the developmental rates varied to reflect the effects of these changes on population processes.

### Influence of the length of developmental diapause

The scenario allowing ticks feeding in the summer of year *y* to undergo a developmental diapause and to begin moulting in the spring of year *y*+1 (Table 1, Theme 2 null hypothesis: “Phase 2 [Mar 21–Jun 21]”, and illustrated in Fig. 2) exhibited a significantly better fit than the scenario assuming a later start of moulting in the summer of year *y*+1 (Table 1, T1.12 vs. T2 ΔDIC: -58.3, Theme 2 alternative hypothesis: “Phase 3 [Jun 22–Sep 22]”). Additionally, the annual peak of questing tick densities predicted by the best-fitting model were, on average, 37% higher for nymphs and 28% higher for adults compared to the model assuming a later start of moulting (Fig. S13), more accurately matching the primarily unimodal questing activity observed in the data.

### Forecasting *I. ricinus* population under climate change scenarios

We chose Murbach site as a representative location for forecasting the *I. ricinus* population for the next 20 years (2023– 2042), considering its higher tick densities compared to the other sites, and assuming that similar climate changes will occur in the Alsace region. Our projections based on the best-fitting model showed that, under moderate to pessimistic scenarios for reducing greenhouse gas emissions, i.e. RCP 4.5, 6.0, and 8.5 (25), the abundance of *I. ricinus* decreased over time (Fig. 6). The extent of the decrease was most noticeable under the RCP 6.0 scenario, with the annual peak of questing tick density decreasing on average by 4.64 per 100m^2^ (95% confidence interval [CI]: 2.77–6.52 per 100m^2^, p-value<0.001) for nymphs and 0.26 per 100m^2^ (95%CI: 0.11–0.40 per 100m^2^, p-value: 0.001) for adults each year (Fig. 6). These decreases corresponded to average reductions of 21.8% and 53.8% in questing nymph and adult densities, respectively, after 20 years, compared to the maximum density values observed at the Murbach site.

**Figure 6.**
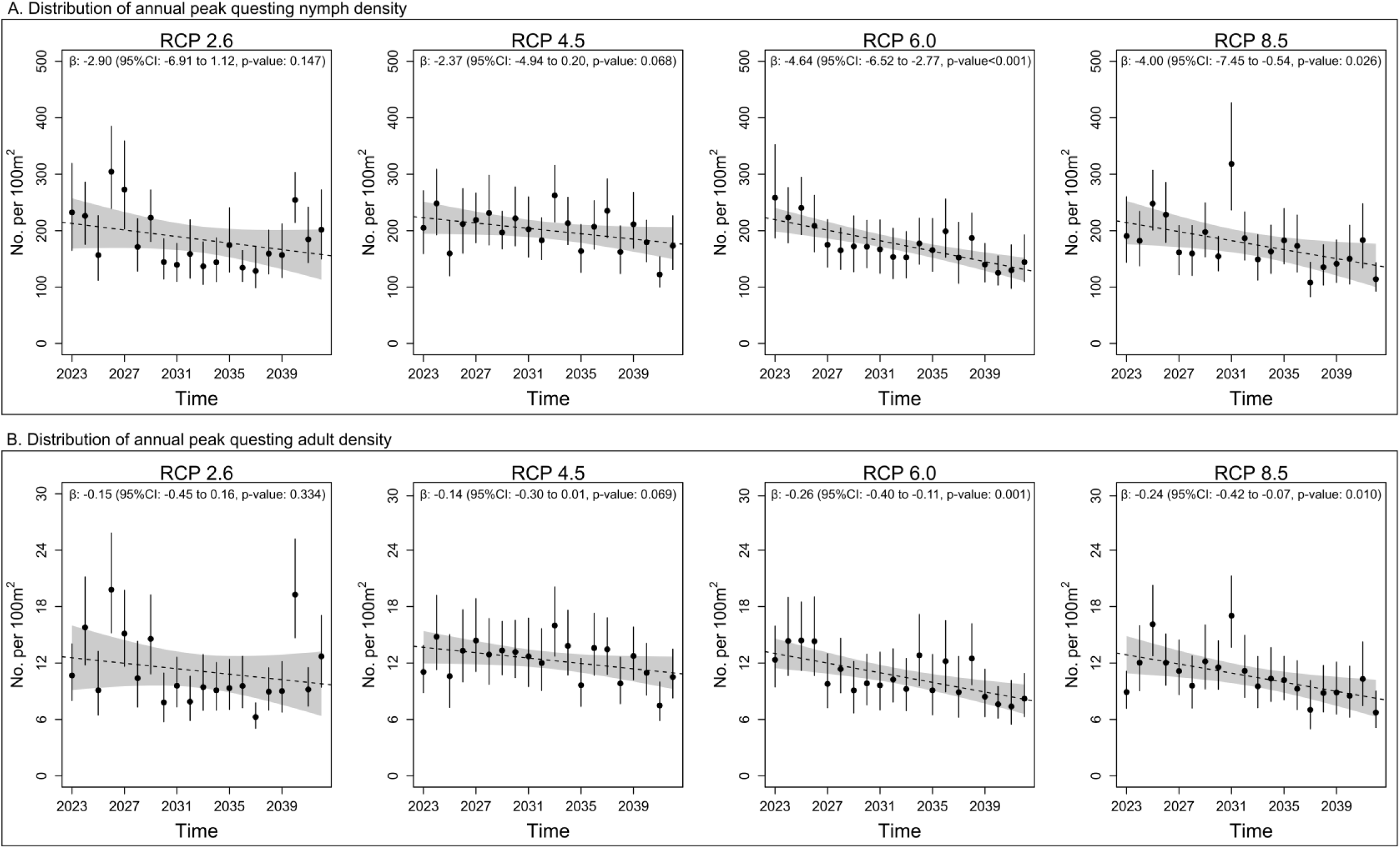
Forecast of questing tick density over the next 20 years until 2042 under Representative Concentration Pathways (RCP). RCP 2.6, 4.5, 6.0, and 8.5 represent climate scenarios from the most optimistic (RCP 2.6) to the most pessimistic (RCP 8.5) view on reducing greenhouse gas emission. Points and vertical lines represent the median and 95^th^ percentile intervals of the annual peak questing (A) nymph and (B) adult densities predicted based on the joint posterior distribution of the best-fitting model and daily mean temperature and saturation deficit values predicted under respective climate scenarios. Linear regression models were fitted to the median values of these annual peak densities, and their mean and 95% confidence interval predictions are illustrated by the dotted lines and shades, respectively, with their results presented on the top of each panel.

## Discussion

Our study quantifies the drivers of *I. ricinus* population dynamics in their natural habitat by fitting a mechanistic model to longitudinal tick density data collected from four sites of ecological and public health relevance. Northeast France and, more broadly, temperate areas of Europe have reported Lyme disease and seen the emergence of tick-borne encephalitis in recent years (22–24), underscoring the importance of understanding the population dynamics of *I.* ricinus, which transmits these diseases. Our model explicitly accounted for the functional relationship between climatic conditions and tick development, the timing of diapauses, and questing and blood-feeding processes. This approach enabled parameter estimation based on field tick density data while addressing its variability (21, 29), allowing us to reproduce the observed population dynamics and, furthermore, generate the first projections of the *I. ricinus* population in one site of the study region under climate change scenarios.

*I. ricinus* has excellent adaptability to a wide range of climate conditions across Europe (1, 18, 19), suggesting that climatic conditions may influence their demographic processes differently to laboratory observations. This was also the case for the studied *I. ricinus* populations. Firstly, the daily moulting rates were estimated to peak at relatively mild temperatures (median: 14.2°C, 95% HDI: 12.5–16.1°C). This finding contrasts with the literature that presented shortening periods between feeding and moulting, and thus increasing moulting rates, with increasing temperature (5, 10). It should be noted that these results from the literature were mainly derived from laboratory experiments where different groups of ticks were exposed to different temperatures under otherwise controlled conditions (5, 10, 17). Importantly, relative humidity was maintained at or near saturation in these experiments, a condition that rarely occurs in natural environments. Indeed, in our study area, relative humidity tended to be low during the hot season in the study region, which may negatively affect the survival of *I. ricinus* undergoing moulting because it temporarily loses its water-absorbing capability in sub-saturated air (4, 30), thereby necessitating adaptation to moult at lower temperatures.

Secondly, allowing ticks feeding in the summer of year y to start moulting only from the spring of year y+1 provided a significantly better explanation for the observed data, compared to allowing a further delayed start in moulting from the summer of year y+1. This could be explained by the observation that the tick density data exhibited a single prominent yearly peak in questing activity, typically during late spring or early summer (19). That is, after a developmental diapause, allowing moulting from the spring, rather than the summer, better prepares a substantial number of unfed ticks to be ready for questing before the main questing season. This dynamic better aligned with the prominent questing peaks, improving the model fit. The alternative scenario of a year-long delay in moulting (i.e. moulting from the summer) might be better suited for *I. ricinus* populations that exhibit a bimodal questing pattern, characterised by an additional, smaller peak in questing activity in the fall (10, 31).

In contrast with the daily moulting rates, our model suggests that the daily oviposition and hatching rates increased with increasing temperature or saturation deficit, consistent with the literature (5, 10). Following peak questing activity in the late spring or early summer, temperatures—and consequently, the drying power of the air, were high when most female adults had their blood meal and were ready for laying eggs. *I. ricinus* is known to continuously lay eggs soon after engorgement unless the temperature drops too low, at which point egg laying may cease (4). Thus, in absence of diapause for engorged female adults and eggs, both oviposition and hatching may have predominantly occurred in the summer. Regarding model fits, saturation deficit explained the observed data significantly better than temperature when it was assumed for both oviposition and hatching processes. However, both climatic variables explained the observed data at similar levels when assumed for either oviposition or hatching. This suggests that variations in saturation deficit were more effective for explaining oviposition and hatching occurring in a synchronised manner, while temperature remained a reliable predictor for each process individually.

Although oviposition and hatching rates were allowed to increase or decrease as functions of environmental variables, we did not explicitly assume a diapause for engorged female adults and eggs due to a lack of data indicating the emergence of eggs and larvae. While the literature suggested that diapause does not seem to occur in engorged *I. ricinus* female adults (4, 32, 33), some studies suggested diapause for eggs (17, 32, 33). Additionally, in our simulation-based model assessments, parameters related to early-stage life forms, i.e. oviposition and hatching, had relatively wide posterior distributions, while a narrower posterior distribution of the hatching-related parameter was observed when the daily oviposition rate was modelled using values derived from controlled laboratory settings. These highlight the importance of, and need for, additional field data on the emergence of larvae to enable further hypothesis testing regarding the presence of diapause and allow for more robust estimation of associated parameters. Such data would include questing larva density data, but this may require more thorough sampling efforts, for example, dragging a larger area, because larvae tend to aggregate more than nymphs and adults, having been hatched from clustered eggs.

We estimated that the daily host contact rates were highest at the Murbach site, suggesting that this study site may have provided a more ideal environment for ticks than the others. Indeed, Murbach has been known to harbour a large abundance of small mammals and, particularly, deer in Northeast France (34, 35). This suggests that the high host abundance in Murbach may have increased the likelihood of encountering a host during questing, leading to more successful progression to the next life stages and thus increased tick abundance. Interestingly, this area has also been associated with high risks of Lyme disease and tick-borne encephalitis (34, 35). This highlights the need for deeper research into the impact of host abundance on tick abundance as well as tick-borne pathogen transmission dynamics.

Climate forecasts over the next 20 years indicated that, under moderate to pessimistic climate scenarios, the abundance of *I. ricinus* at the Murbach site decreases significantly. This highlights the need for continued monitoring of the status of *I. ricinus* populations in the study region, as well as their role as reservoir and vector of Lyme disease, tick-borne encephalitis, or other zoonotic pathogens. In particular, our forecasts provide insights into the impact of temperature increases in areas where *I. ricinus* has already been established under similar climatic conditions. This needs to be considered along with the geographical expansion of *I. ricinus* towards regions at higher latitudes and higher elevations previously uninhabited by *I. ricinus* (36–38), when assessing the impact of climate change on the overall disease burden by *I. ricinus*-borne pathogens. However, caution must be exercised in interpreting this result because climatic changes would also affect tick mortality, host population dynamics and patterns of land-use, factors which were not explicitly addressed by our model. Additionally, considering ticks’ adaptability evidenced in various parts of Europe (18, 19), tick development may exhibit different functional relationships with future climatic conditions compared with those estimated based on the observed data.

Further points merit consideration when interpreting the study results. Firstly, our model assumed that baseline daily mortality and host contact rates remained constant throughout each year due to the lack of available information. These rates are likely to vary in response to changes in energy reserves, climatic conditions, and host availability (31, 39), possibly leading to yearly variations in tick abundance, as indicated by variations in questing tick density in this study. Secondly, although our model structure allowed for assessing the timing of diapause (spring vs. summer), it was constrained by the way calendar years were divided, which may not fully reflect the field reality. This is because, although the division was made based on changes in day length, which is a primary determinant of diapause (33, 39), the initiation and, particularly, termination of diapause could also be influenced by other factors, such as local climatic conditions and heterogeneities in the behaviour of individual ticks (33, 39). Thirdly, the functional relationship between climatic conditions and tick development were constrained by the fixed variance of the Gamma distributions assumed to underlie those relationships, although sensitivity analysis did not indicate major changes in the study results. These points underline the need for further demographic data to improve model inference, such as temporal variations in mortality and host abundances, as well as the emergence of newly moulted ticks or laid eggs.

In conclusion, our study elucidated the environmental and ecological drivers of *I. ricinus* population dynamics in the Alsace region of Northeast France. While oviposition and hatching processes demonstrated similar dynamics to those suggested by laboratory experiments and field observations in other regions, moulting exhibited unique dynamics in its relationship with climatic conditions and delays through developmental diapause. These characteristics were considered to drive the predominantly unimodal questing pattern observed in the study region. Our findings emphasise the importance of further research tailored to various regions where *I. ricinus* has established, to assess heterogeneities in the mechanisms behind *I. ricinus* population dynamics. Furthermore, our modelling framework can also be expanded to other *Ixodidae* ticks of public and animal health significance. Finally, when combined with relevant epidemiological data, our modelling framework will allow the assessment of the transmission of *Ixodes*-borne diseases, such as Lyme disease and tick-borne encephalitis.

## Materials and Methods

### Longitudinal questing tick density and climate data

Between 2013 and 2022, we collected host-seeking *I. ricinus* ticks (questing nymphs and adults) at four sites (Dannemarie, Illkirch, Murbach, and Niedermunster) in the Alsace region of Northeast France. Climate data were based on daily mean temperature and relative humidity near the surface from the Copernicus Climate Change Service (25, 40, 41). See ‘Tick sampling and climate data’ in Appendix S1 Methods and Results for details.

### Mechanistic model for tick population dynamics

#### Model structure

Figures 1 and 2 illustrate the overall model structure. Table S1 describes input parameters, and Table 2 details the estimated parameters. Hereinafter, unfed ticks (L_u_, N_u_, and A_u_) defined ticks that have newly moulted or hatched but have not yet begun seeking a host, and questing ticks (L_q_, N_q_, and A_q_) were ticks actively seeking a host. Engorged ticks (L_e_, N_e_, and A_e_) defined ticks that have fed but have not yet progressed to the next life stage.

The progression of *I. ricinus* occurs through development (oviposition, hatching, larva-to-nymph moulting, and nymph-to-adult moulting), questing, host contact, and mortality. Of these processes, development and questing are influenced by meteorological conditions and the occurrence of developmental and behavioural diapauses (33, 39). Our model accounted for these meteorological impacts by allowing ticks to advance through life stages at developmental and questing rates changing daily according to climatic variations. Additionally, our model accounted for developmental diapause (delays in moulting) and behavioural diapause (delays in questing) by allowing for moulting based on the timing of feeding, and for questing based on the timing of moulting, across four distinct phases of the year. Phase 1 (Jan 1–Mar 20) and Phase 4 (Sep 23–Dec 31) were periods when moulting processes were halted because of short daylengths (day is longer than night) and low temperatures during these months (33, 39). In contrast, Phase 2 (Mar 21–Jun 21) and Phase 3 (Jun 22–Sep 22) were periods where moulting processes could occur, divided at summer solstice when day length starts to decrease. Following this calendar, engorged larvae and nymphs were categorised into the following compartments according to the timing of feeding in the year: ticks fed in Phases 1 and 2 (L_e1·2_ and N_e1·2_), and those fed in Phases 3 and 4 (L_e3·4_ and N_e3·4_). Questing activity was allowed throughout the four phases, with its daily rate being influenced by meteorological conditions, but depending on the timing of moulting. Thus, to account for behavioural diapause, the compartments for unfed nymphs (N_u_) and adults (A_u_) were divided based on the timing of moulting: ticks moulted in Phase 2 (N_u2_ and A_u2_), and those moulted in Phase 3 (N_u3_ and A_u3_) (see *Developmental rate* and *Questing rate* below for details).

The following sections detail how developmental, questing, and moulting, and mortality rates were modelled.

#### Developmental rate

The daily oviposition, hatching, and larva-to-nymph and nymph-to-adult moulting rates were modelled to be associated with climatic conditions under Theme 1 hypotheses. Theme 1 null hypothesis assumed that these developmental rates continued to increase with daily mean temperature, following the inverse of the power-law curves estimated by Ogden, Lindsay (5) (Table S1, η_𝑖=1,…,4_), derived from the observed number of days required for *Ixodes scapularis* to progress to a subsequent life stage under different temperature conditions in a laboratory setting. According to these curves, peak rates occurred exclusively at the maximum temperature observed.

In contrast, Theme 1 alternative hypothesis allowed that these developmental rates increased, peaked, and decreased across the range of a given climatic condition (temperature or saturation deficit) observed in the ticks’ natural habitats, following the shape of a Gamma distribution (Table 2, γ_𝑖=1,…,4_). For example, the daily rate of developmental process 𝑖 ( 𝛾_𝑖𝑡_) was hypothesised to vary with daily mean temperature *t*, following the shape of a Gamma distribution defined by its mode (θ_𝑖_) and variance (ε_𝑖_):

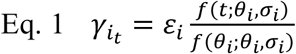

Here, 𝑓(𝑡; 𝜃_𝑖_, 𝜎_𝑖_) represented the probability density of a Gamma distribution at temperature 𝑡, with 𝜃_𝑖_ and 𝜎_𝑖_ being its mode and standard deviation, respectively, derived from the shape and rate parameters of a Gamma distribution. This probability density was normalised by the maximum probability density, 𝑓(𝜃_𝑖_; 𝜃_𝑖_, 𝜎_𝑖_), and then converted into a rate using a scaling parameter 𝜀_𝑖_ . Thus, 𝜀_𝑖_ represented the peak rate of developmental process 𝑖 occurring at a temperature of 𝜃_𝑖_. Under this hypothesis, 𝜃_𝑖_ and 𝜀_𝑖_ were estimated through model fitting, while 𝜎_𝑖_ was fixed to a value that covered a reasonably wide temperature range within which ticks were considered to be active (Tables 2 and S1). We assumed that larva-to-nymph and nymph-to-adult moulting rates peaked under the same climatic condition (𝜃_3_ = 𝜃_4_), but at different levels of intensity (𝜀_3_ ≠ 𝜀_4_).

The timing of developmental diapause (Theme 2) was assessed as follows. The null hypothesis assumed that ticks that fed during Phase 2 (when day length increases, Mar 21–Jun 21) moulted in Phase 3 of the same year (dark brown shades in Fig. 2) or Phase 2 of the following year (pale brown shades in Fig. 2), based on climatic conditions (see Theme 1). Ticks that fed during Phase 3 (when day length decreases, Jun 22–Sep 22) moulted in Phase 2 or Phase 3 of the following year (pale brown shades in Fig. 2), still based on climatic conditions (see Theme 1). In Theme 2 alternative hypothesis, these ticks were allowed to moult only from Phase 3 of the following year, whilst the same assumption was made for ticks that fed during Phase 2.

#### Questing rate

The daily questing rate (𝜏_𝑙,𝑠,𝑑_) was defined as the rate at which unfed ticks are active and looking for a host, and modelled as follows. The predicted questing tick density from a generalised additive model (GAM fit values) was assumed to reflect trends in daily questing rates. The GAM fitted values were converted into a daily questing rate multiplied by a scaling parameter δ, which was estimated by fitting the mechanistic model (see ‘Generalised additive model (GAM) analysis and results’ in Appendix S1 Methods and Results and Figs S14).

In addition, our model accounted for behavioural diapause (delays in questing) by allowing ticks to start questing based on the timing of their moulting. Firstly, ticks that moulted in Phase 3 were modelled to start questing from Phase 2 of the following year (pale green shades in Fig. 2). In contrast, those that moulted in Phase 2 were allowed to begin questing without a behavioural diapause (dark green shades in Fig. 2).

#### Host contact rate

Questing ticks were modelled to take a blood meal following a daily host contact rate, 𝛽_𝑙,𝑠_, as follows:

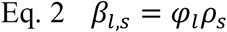

where 𝛽_𝑙,𝑠_was assumed to vary between the life stages (subscript 𝑙) and across the four study sites (subscript 𝑠). The baseline host contact rates ( 𝜑_𝑙_) was assumed different for larvae, nymphs, and adults, but with the same ratios as estimated by Dobson, Finnie and Randolph (8), and were scaled differently (𝜌_𝑠_) across the four sampling sites, to account for the between-site variability resulting from tick and host densities and host species. β was assumed constant over time, with the premise that temporal changes in the daily questing rate would explain temporal changes in the probability of feeding on a host.

#### Mortality rate

The daily mortality rates were based on values derived in the literature and assumed to increase linearly with tick population size (8, 27) (see ‘Mechanistic model parameters’ in Appendix S1 Methods and Results for details).

### Model fitting and hypothesis testing

Model fitting was conducted within an HMC Bayesian inference framework using the *RStan* package (42) in R version 4.3.2 (43). For the likelihood function, questing tick density was assumed to follow a negative binomial distribution (see ‘Likelihood function for mechanistic model fitting’ in Appendix S1 Methods and Results for details).

First, we assessed the model’s ability to recover true parameter values by fitting it to synthetic data (see ‘Simulation-based model assessments and results’ in Appendix S1 Methods and Results). Secondly, we estimated parameters related to tick demographic processes by fitting the model to the observed longitudinal questing tick density data.

Model fits under different assumptions on the functional relationships between climatic conditions and tick development (Theme 1) were assessed based on the deviance information criterion (DIC) as follows. The baseline model (Table 1, T0) assumed that (i) all types of developmental rates increased with temperature following the inverse of the power-law curves estimated by Ogden, Lindsay (5) (Theme 1 null hypothesis) and (ii) ticks that fed a blood meal in Phase 3 moulted from Phase 2 of the following year after undergoing a developmental diapause (Theme 2 null hypothesis).

During model fit comparison, one specific type of developmental rate was assumed to vary with temperature or saturation deficit, following a Gamma distribution shape (Table 1, T1.1–T1.6), in contrast to the power-law function assumed in the baseline model. If this alternative assumption resulted in a decrease in DIC greater than 5 units, it was considered to indicate a significantly better explanation of the observed data and replaced the null hypothesis assumed in the baseline model. Subsequently, other types of developmental rate were assessed in the same manner (Table 1, T1.7 – T1.12), and the best-fitting mechanistic model—a model with a set of hypotheses that best explains the questing tick density data—was determined when no alternative hypotheses led to a DIC reduction greater than 5 units.

As a posterior predictive check, the densities of questing nymphs and adults were simulated based on the joint posterior distribution of the best-fitting mechanistic model and compared with the observed densities. For a given developmental process 𝑖, its functional relationship with the climatic variable selected in the best-fitting mechanistic model was described based on 𝜃_𝑖_ and 𝜀_𝑖_. If daily mean saturation deficit was selected in the best-fitting model, the functional relationship with daily mean temperature was also explored in a separate model fitting to help the interpretation of study results. Finally, to explore how the length of developmental diapause on a model fit (Theme 2), we compared the best-fitting model to the scenario that allowed extended delays in moulting for ticks that fed in Phase 3 (Table 1, T2).

Sensitivity analyses of the assumed daily baseline mortality and host contact rates were conducted to assess their impacts on the functional relationships between developmental rates and climatic conditions. Specifically, for each analysis, the relationships were evaluated using the joint posterior distribution, with one of the baseline mortality or host contact rates varied by ±20%.

### Projections under different climatic scenarios

We forecast the *I. ricinus* population at the Murbach site until 2042 based on the climate changes predicted under the Representative Concentration Pathways (RCP) 2.6, 4.5, 6.0, and 8.5 scenarios (44)—corresponding to mild to severe impacts of greenhouse gas emission—and the parameter estimates from the best-fitting mechanistic model. The *I. ricinus* population at the Murbach site was chosen as it exhibited highest tick densities in the observed data. For each RCP scenario, forecasting was performed by simulating the *I. ricinus* population for each set of joint posterior parameter estimates, based on the predicted changes in the climatic variable selected in the best-fitting model. For each iteration, we recorded the annual peak questing tick densities as an indicator of tick abundance. After the simulation, we computed their median values for each year and then fitted a linear regression model to these median values over the forecasted years.

## Acknowledgements

This research is supported by French Agence Nationale de la Recherche (MozArt project, project number ANR-22-CE35-0003, www.mozartonehealth.com). National Reference Center for Borrelia is supported by Santé Publique France. WJE and JV were funded by the National Institute for Health Research (NIHR) Health Protection Research Unit in Modelling and Health Economics (grant code 200908). The views expressed are those of the author(s) and not necessarily those of the NIHR, UK Health Security Agency or the Department of Health and Social Care. We gratefully thank Lisa Baldinger and Marine Engel for collecting the ticks, and for Laure Bournez for discussions on the biology of Ixodes ricinus. Finally, we acknowledge the World Climate Research Programme’s Working Group on Coupled Modelling, which is responsible for CMIP, and we thank the climate modelling groups (listed in Table 2 of this paper) for producing and making available their model output. For CMIP the U.S. Department of Energy’s Program for Climate Model Diagnosis and Intercomparison provides coordinating support and led development of software infrastructure in partnership with the Global Organization for Earth System Science Portals.

---

Figure S1. Description of longitudinal tick density data

Figure S2. Model assessments using synthetic tick density data

Figure S3. Difference between the predicted and observed questing nymph densities

Figure S4. Difference between the predicted and observed questing adult densities

Figure S5. Parameter estimates from the best-fitting model

Figure S6. Impact of varying daily egg mortality rates

Figure S7. Impact of varying daily larva mortality rates

Figure S8. Impact of varying daily nymph mortality rates

Figure S9. Impact of varying daily adult mortality rates

Figure S10. Impact of varying baseline daily host contact rates of larvae

Figure S11. Impact of varying baseline daily host contact rates of nymphs

Figure S12. Impact of varying baseline daily host contact rates of adults

Figure S13. Comparison of annual peak questing tick densities predicted under Theme 2 null and alternative hypotheses

Figure S14. Effects on questing Ixodes ricinus density estimated by the best-fitting generalised additive models (GAMs)

Table S1. Input parameters for the mechanistic model. Notations, definitions, values and sources or assumptions.

Table S2. Results of the best-fitting generalised additive models (GAMs)

Appendix S1. Methods and Results

