## Supplementary Information for "Environmental drivers of *Ixodes ricinus* tick population dynamics: mechanistic modelling using longitudinal field surveys and climate data"

### Supplementary Figures and Table

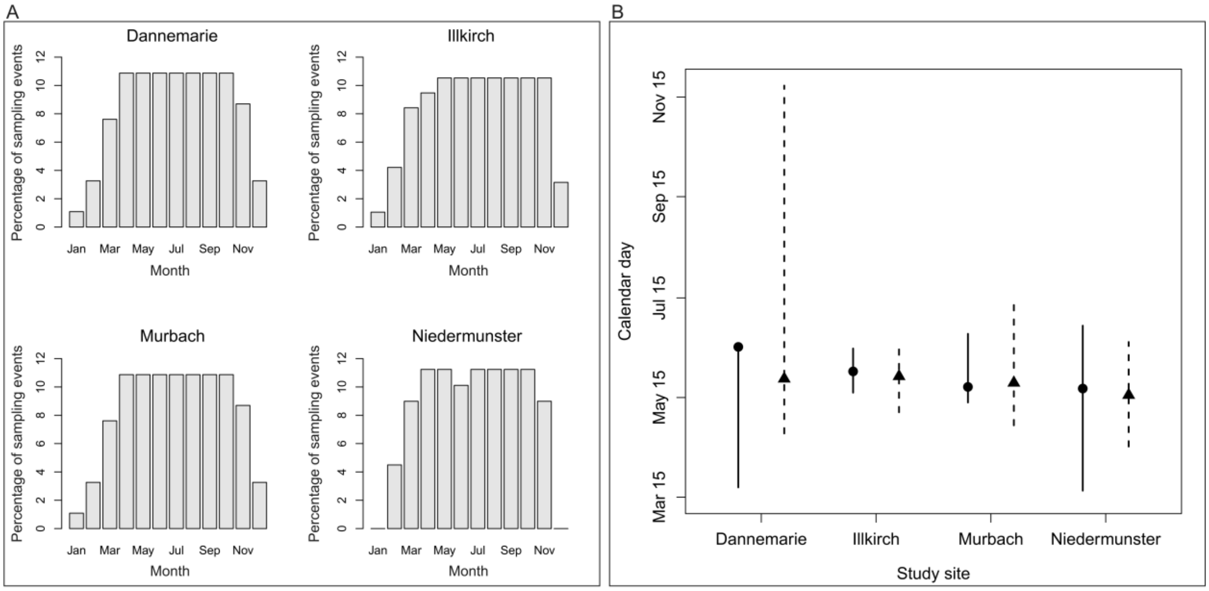

**Figure S1. Description of longitudinal tick density data. (A)** Distribution of sampling sessions by month and sampling site across 2013–2022. **(B)** Distribution of calendar days with yearly peak questing tick density. The points and vertical lines represent the median and 95% percentile intervals, respectively, of calendar days on which the highest questing nymph (circles and solid lines) or adult (triangles and dashed lines) density (number per $100m^2$) was observed each year.

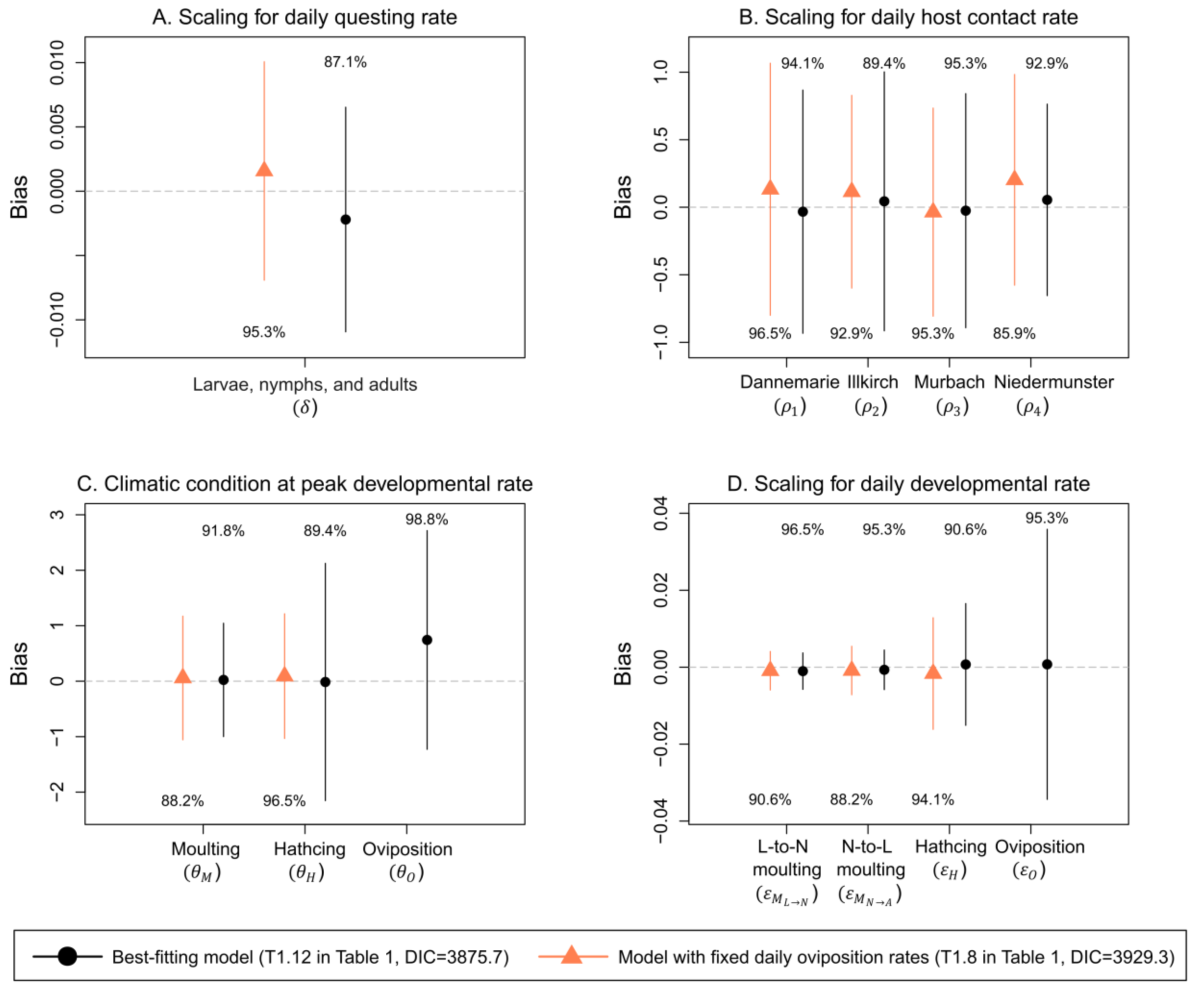

**Figure S2. Model assessments using synthetic tick density data.** Points and vertical lines represent the mean and ±1 standard deviation of the differences between the true parameter values (used to generate synthetic data) and the posterior modes (obtained from model fitting to the synthetic data) from 85 simulations. Circles and triangles represent the results from the best-fitting model and the model in which the daily oviposition rate was modelled using values from controlled laboratory settings. Numbers show the percentage of simulations where each true parameter value fell within the 95% highest density interval of the corresponding posterior distribution.

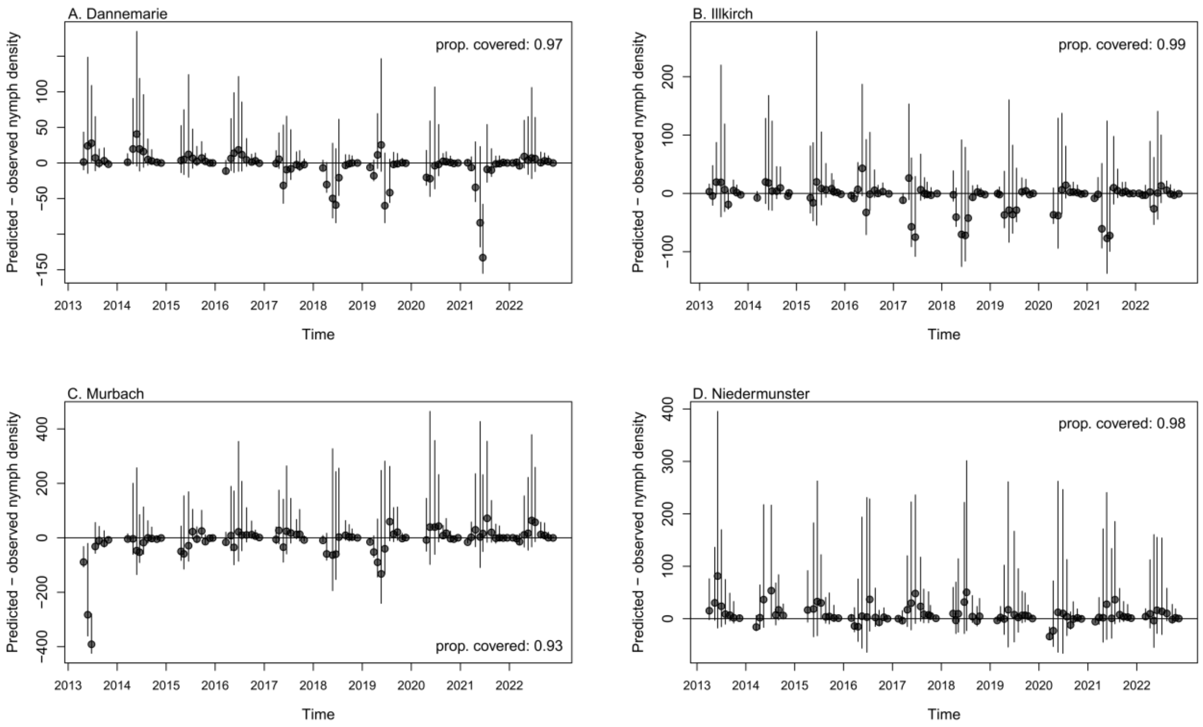

**Figure S3. Difference between the predicted and observed questing nymph densities.** For each sampling site (A–D) and date (x-axis), the observed density was subtracted from the 95th percentile interval of the predicted density (y-axis). The predicted densities were estimated based on the joint posterior distribution of the best-fitting model. "Prop. Covered" corresponds to the proportion of sampling sessions for which the observed density was within the 95th percentile interval of the predicted density.

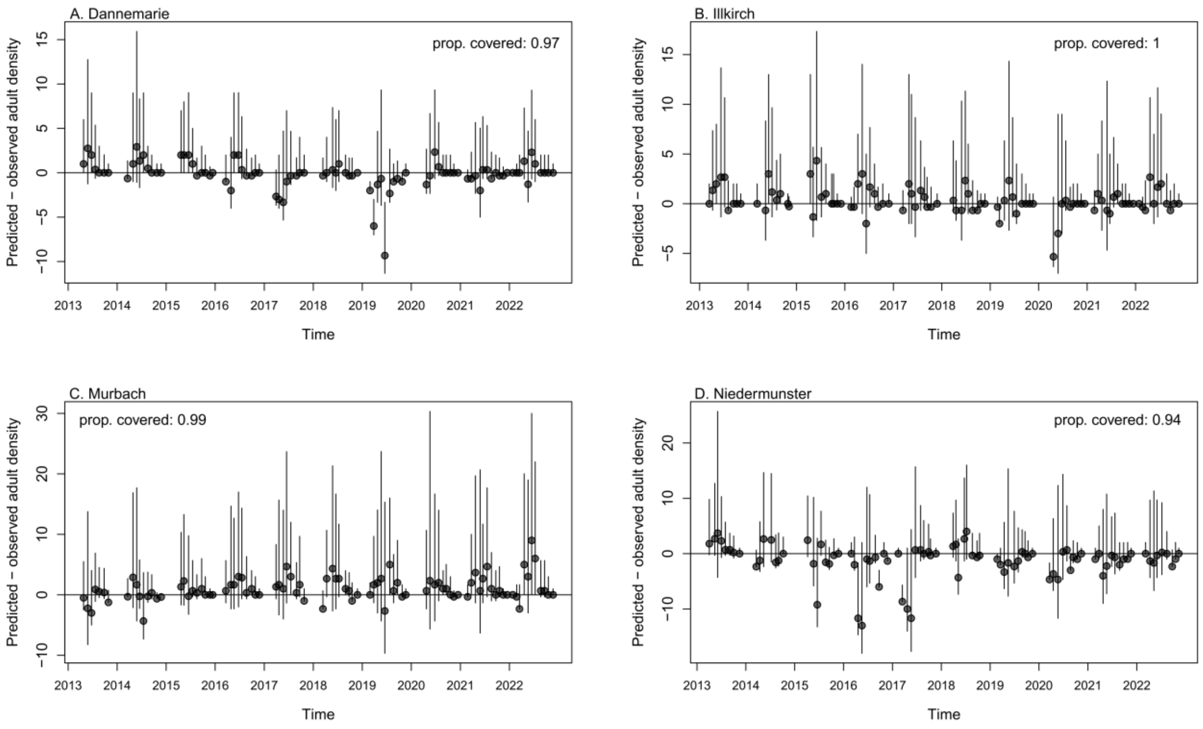

**Figure S4. Difference between the predicted and observed questing adult densities.** For each sampling site (A–D) and date (x-axis), the observed density was subtracted from the 95th percentile interval of the predicted density (y-axis). The predicted densities were estimated based on the joint posterior distribution of the best-fitting model. "Prop. Covered" corresponds to the proportion of sampling sessions for which the observed density was within the 95th percentile interval of the predicted density.

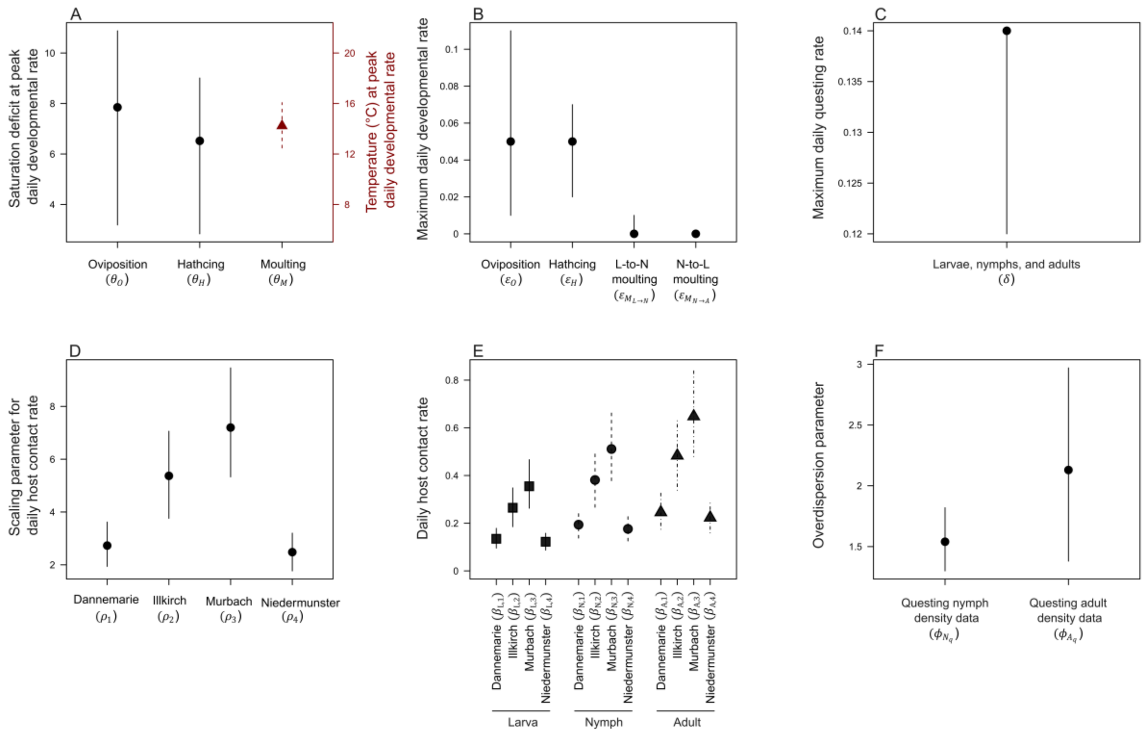

**Figure S5. Parameter estimates from the best-fitting model.** See Table 2 for the details of the parameters estimated.

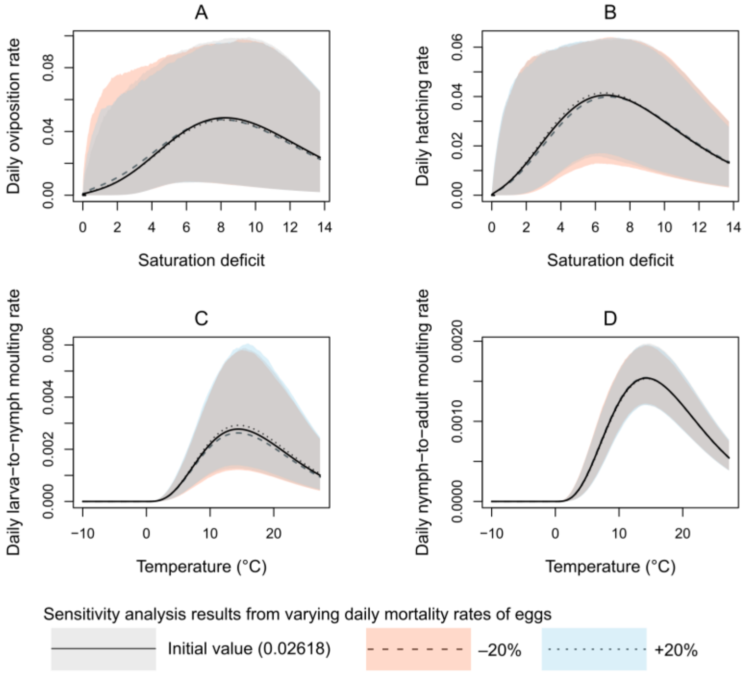

**Figure S6. Impact of varying daily egg mortality rates.** The solid lines and grey shades represent the median and 95th percentile intervals of the developmental rates estimated using the initial mortality rate. The dashed lines and reddish shades represent estimates based on a mortality rate 20% lower, while the dotted lines and bluish shades represent estimates based on a mortality rate 20% higher.

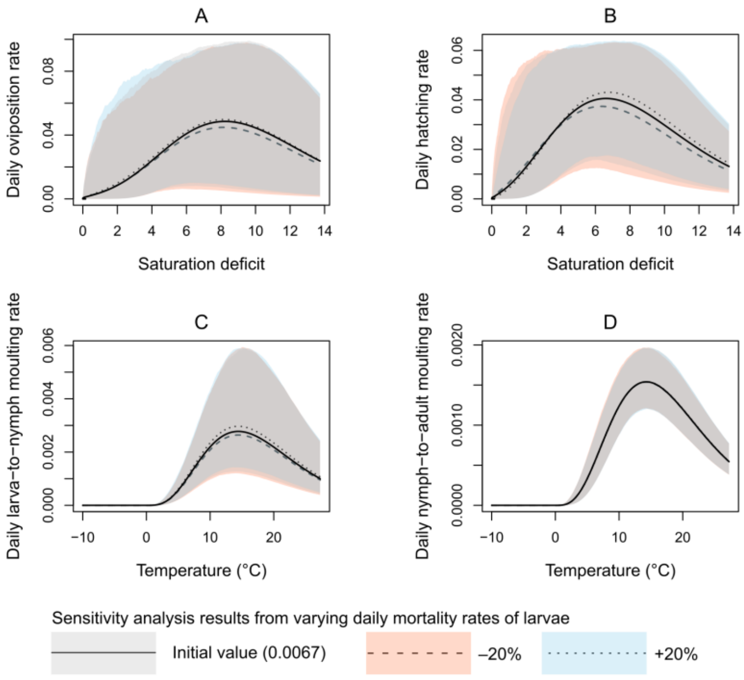

**Figure S7. Impact of varying daily larva mortality rates.** The solid lines and grey shades represent the median and 95th percentile intervals of the developmental rates estimated using the initial mortality rate. The dashed lines and reddish shades represent estimates based on a mortality rate 20% lower, while the dotted lines and bluish shades represent estimates based on a mortality rate 20% higher.

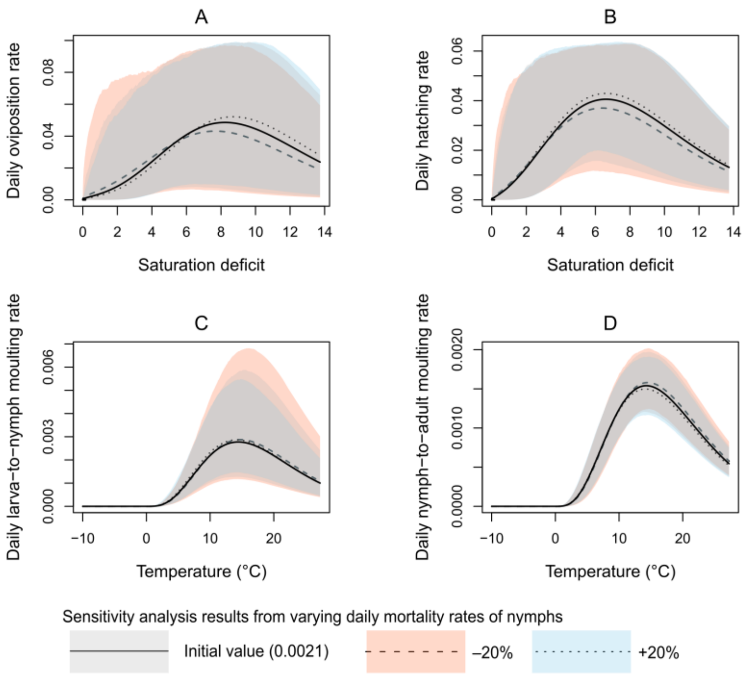

**Figure S8. Impact of varying daily nymph mortality rates.** The solid lines and grey shades represent the median and 95th percentile intervals of the developmental rates estimated using the initial mortality rate. The dashed lines and reddish shades represent estimates based on a mortality rate 20% lower, while the dotted lines and bluish shades represent estimates based on a mortality rate 20% higher.

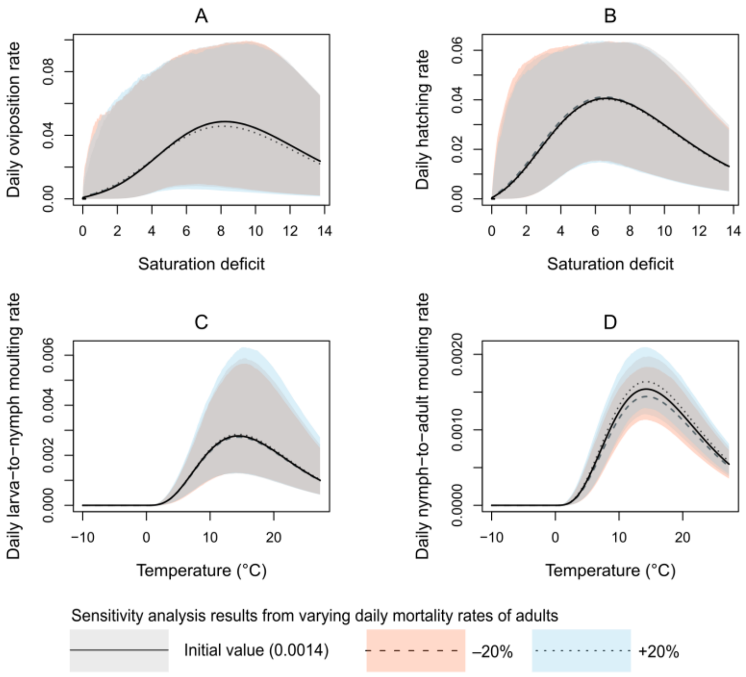

**Figure S9. Impact of varying daily adult mortality rates.** The solid lines and grey shades represent the median and 95th percentile intervals of the developmental rates estimated using the initial mortality rate. The dashed lines and reddish shades represent estimates based on a mortality rate 20% lower, while the dotted lines and bluish shades represent estimates based on a mortality rate 20% higher.

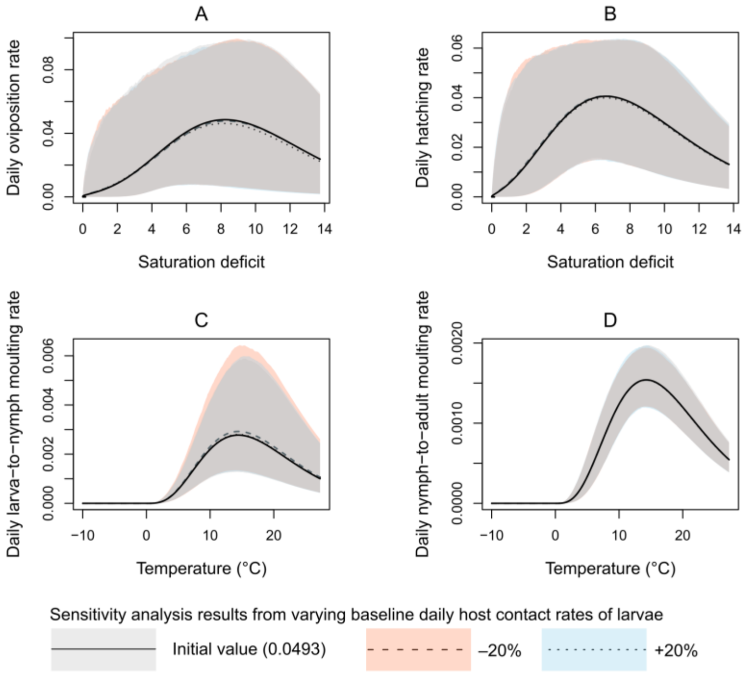

**Figure S10. Impact of varying baseline daily host contact rates of larvae.** The solid lines and grey shades represent the median and 95th percentile intervals of the developmental rates estimated using the initial baseline host contact rate. The dashed lines and reddish shades represent estimates based on a baseline host contact rate 20% lower, while the dotted lines and bluish shades represent estimates based on a baseline host contact 20% higher.

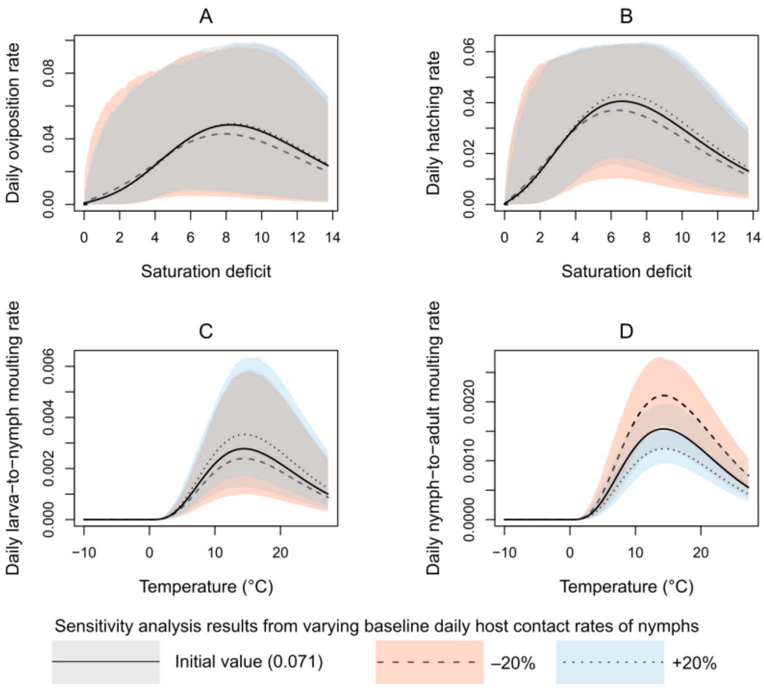

**Figure S11. Impact of varying baseline daily host contact rates of nymphs.** The solid lines and grey shades represent the median and 95th percentile intervals of the developmental rates estimated using the initial baseline host contact rate. The dashed lines and reddish shades represent estimates based on a baseline host contact rate 20% lower, while the dotted lines and bluish shades represent estimates based on a baseline host contact 20% higher.

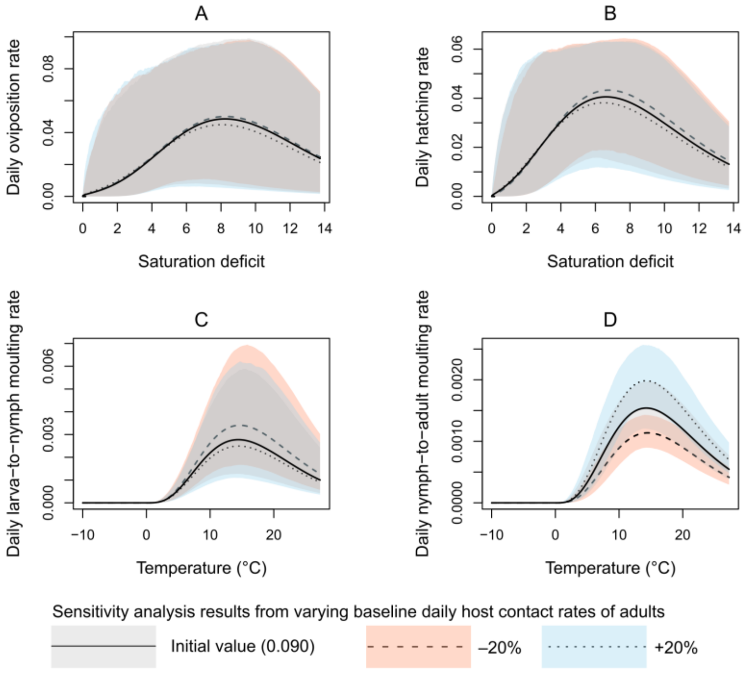

**Figure S12. Impact of varying baseline daily host contact rates of adults.** The solid lines and grey shades represent the median and 95th percentile intervals of the developmental rates estimated using the initial baseline host contact rate. The dashed lines and reddish shades represent estimates based on a baseline host contact rate 20% lower, while the dotted lines and bluish shades represent estimates based on a baseline host contact 20% higher.

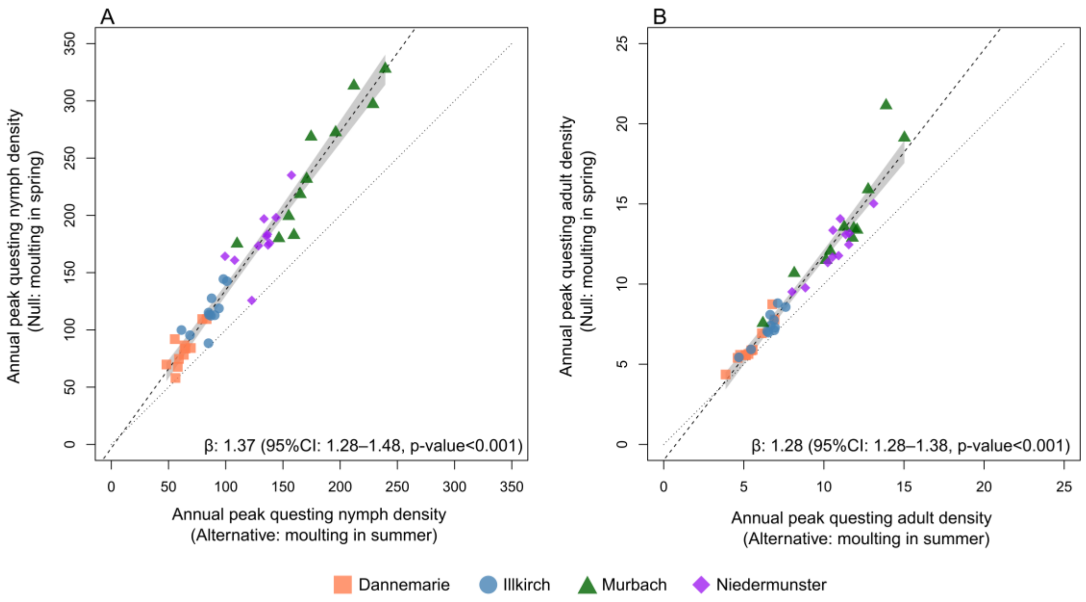

**Figure S13. Comparison of annual peak questing tick densities predicted under Theme 2 null and alternative hypotheses.** The points represent the median values of the annual peak questing nymph (A) and adult (B) densities predicted by, for ticks that fed in summer, assuming moulting from the following spring (Theme 2 null hypothesis, y-axis) or assuming moulting from the following summer (Theme 2 alternative hypothesis, x-axis). Different point shapes represent different study sites. The dashed lines and grey shades represent the linear regression lines and its 95% confidence intervals fitted to those points.

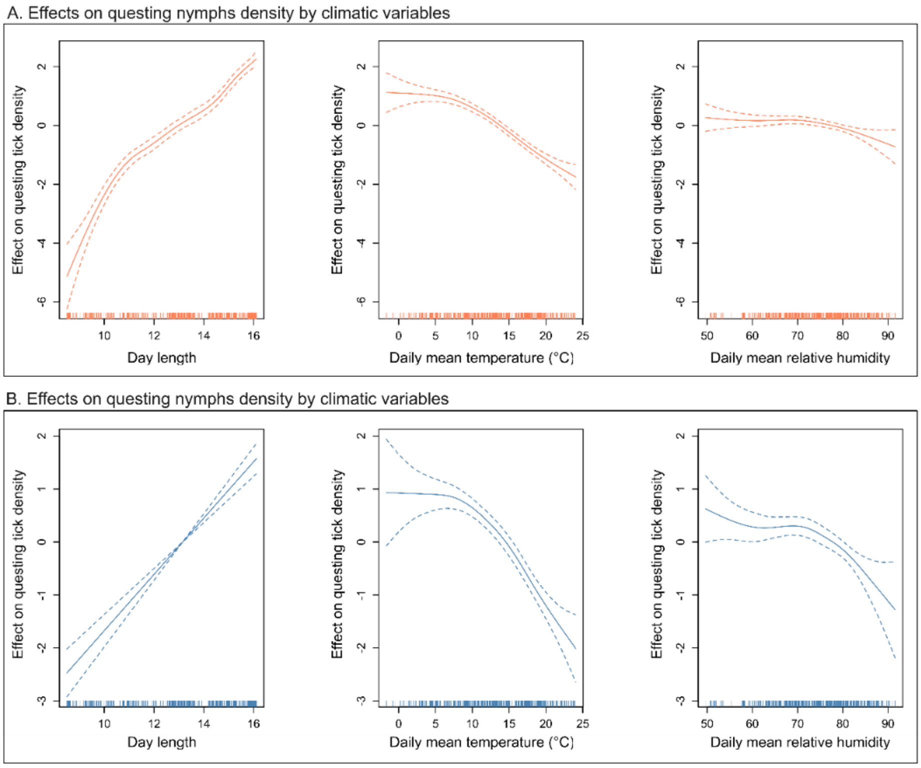

**Figure S14. Effects on questing Ixodes ricinus density estimated by the best-fitting generalised additive models (GAMs).** Panels A and B display the effects of a given climatic variable on questing nymph (A) and adult (B) densities, respectively. The vertical lines on the x-axis indicated sampling sessions that occurred at the corresponding climatic conditions.

Table S1. Input parameters for the mechanistic model. Notations, definitions, values and sources or assumptions.

| Notation | Definition | Value | Source or assumption |
|---|---|---|---|
| $n_{new\ eggs}$ | Number of eggs laid by one female adult | 1,597 | Macleod (1), Sonenshine (2) |
| $m_{0_l}$ | Daily background mortality for unfed and questing ticks by life stage [a] | 0.02618 (egg) 0.0067 (larva), 0.0021 (nymph), 0.0014 (adult) | Dobson, Finnie and Randolph (3); Daniel, Cerný (4) |
| $m_2$ | Daily background mortality for engorged ticks | 0.00274 | Engorged ticks survive for one year if not moulting or laying eggs (5, 6). |
| $\varphi_l$ | Baseline daily host contact rate by life stage | 0.0493 (larva), 0.071 (nymph), 0.09 (adult) | Dobson, Finnie and Randolph (3) |
| $\sigma$ | The standard deviation of the Gamma distribution, which specified changes in the developmental rates by climate conditions | Temperature: 8°C<br>Saturation deficit: 4 | The standard deviation was set to encompass reasonably wide ranges of either temperature or saturation deficit values. |
| $GAM\ fit_{l,s,d}$ | Temporal trends in daily question rates by life stage and site | see 'Generalised additive model (GAM) analysis and results' in SI | Temporal trends in daily questing rates were assumed to follow daily questing tick density predicted by the best-fitting generalised additive model (GAM). |
| $GAM\ fit_{l_{max}}$ | The maximum predicted tick density by life stage | see 'Generalised additive model (GAM) analysis and results' in SI | |
| $\eta_{O_t}$ | Daily oviposition rate at temperature *t* under Theme 1 null hypothesis | $1/(1300t^{-1.42})$ | Ogden, Lindsay (7). Temperatures below zero were assumed zero when daily developmental rates were determined by these functions. |
| $\eta_{H_t}$ | Daily hatching rate at temperature *t* under Theme 1 null hypothesis | $1/(34234t^{-2.27})$ | |
| $\eta_{M_{L\to N_t}}$ | Daily larva-to-nymph moulting rate at temperature *t* under Theme 1 null hypothesis | $1/(101181t^{-2.55})$ | |
| $\eta_{M_{N\to A_t}}$ | Daily nymph-to-larva moulting rate at temperature *t* under Theme 1 null hypothesis | $1/(1596t^{-1.21})$ | |

| **Climate condition (2008–2022)** [b] | **median** | **95th percentile interval** | |
|---|---|---|---|
| Daily mean 2m temperature [c] | 10.2°C | -10.0–27.3°C | Daily statistics calculated from ERA5 data. |
| Daily mean relative humidity [c] | 77.1% | 42.1–97.3% | Daily statistics calculated from ERA5 data. |
| Daily mean saturation deficit | 2.2 | 0.15–13.7 | Computed based on ERA 5 daily mean temperature and relative humidity using the equation in Randolph and Storey (8) |
| Day length | 12.3 hrs | 8.3–12.3 hrs | The data were extracted for each day and each site using the *daylength* function of the *geosphere* package. |

a The overall mortality, $m_1$, was assumed to increase with population size from these daily background mortality rates.
b The climate conditions were smoothed with a Gaussian kernel of 21 days to account for their cumulative impact on tick host demography processes.
c Daily mean 2m temperature and relative humidity data under climate change scenarios between 2023 and 2042 were sourced from the Copernicus Climate Change Service under the following settings. Experiment: RCP 2.6, 4.5, 6.0, and 8.5, Model: IPSL-CM5A-LR (IPSL, France), Ensemble number: r1i1p1

Table S2. Results of the best-fitting generalised additive models (GAMs)

| Parameter | Model outputs | | |
|---|---|---|---|
| | Effective degrees of freedom | $\chi^2$ statistics | p-value |
| **Best-fitting GAM for questing nymph density** | | | |
| **Random-effect terms** | | | |
| **Sampling site** | 2.886 | 76.39 | <0.001 |
| **Sampling year** | 7.145 | 35.27 | <0.001 |
| **Cubic spline terms (maximum k=10)** [a] | | | |
| **Day length** | 5.149 | 505.78 | <0.001 |
| **Daily mean temperature** | 3.430 | 203.04 | <0.001 |
| **Daily mean relative humidity** | 2.709 | 10.68 | 0.02 |
| **Deviance explained** | 75.3% | | |
| **Best-fitting GAM for questing adult density** | | | |
| **Random-effect terms** | | | |
| **Sampling site** | 2.881 | 94.13 | <0.001 |
| **Sampling year** | 6.287 | 21.54 | <0.001 |
| **Cubic spline terms (maximum k=10)** [a] | | | |
| **Day length** | 1.001 | 120.17 | <0.001 |
| **Daily mean temperature** | 3.241 | 110.58 | <0.001 |
| **Daily mean relative humidity** | 3.150 | 19.80 | <0.001 |
| **Deviance explained** | 57.3% | | |

a k is the number of basis functions for cubic splines, and its maximum value was set considering the annual number of sampling sessions (9 – 11 sessions per year).

### Appendix S1 Methods and Results

#### Tick sampling and climate data

Sampling sites included Dannemarie (47°41'56.7"N 7°09'15.1"E), Illkirch (48°31'06.3"N 7°44'35.9"E), Murbach (47°55'18.2"N 7°09'21.3"E), and Niedermunster (48°26'02.6"N 7°24'28.6"E). At each visit, the ticks were collected from the vegetation and litter by dragging a $1m^2$ piece of white cotton sheet over an area of $10m^2$ and inspecting the cloth for ticks every $10m^2$; at each visit, a total of 30 draggings were performed in order to sample a total vegetation area of $300m^2$. The same area was collected at each visit. The captured nymph and adult ticks were removed from the cloth, identified, and counted, and the density per $100m^2$ was calculated. At least two collectors participated in each visit, which occurred every month when the weather conditions were favourable for sampling.

At each visit, the ground temperature and relative humidity were measured and recorded with an LCD digital hygro-thermometer. Additionally, for each sampling site, observed and projected daily mean temperature and relative humidity near the surface were sourced from the Copernicus Climate Change Service for the period 2008–2022 (9, 10) and for the following 20 years under different Representative Concentration Pathway (RCP) scenarios (11), including RCP 2.6, 4.5, 6.0, and 8.5. A higher RCP value indicated a higher level of greenhouse gas emissions, and therefore, global surface temperatures, over time.

Daily mean temperature and relative humidity from the Copernicus Climate Change Service were highly correlated with those recorded from the field. Therefore, the metrics from the Copernicus Climate Change Service were used for fitting models and projecting *I. ricinus* tick abundance under the RCP scenarios. Additionally, these metrics were used to calculate daily mean saturation deficit, based on the formula provided by Randolph and Storey (8). Day length was obtained using the *daylength* function of the *geosphere* package (12) in R. 4.3.2 (13). Climate data were smoothed with a Gaussian kernel of 21 days to account for their cumulative impact on tick host demography processes.

#### Simulation-based model assessments and results

Models were fitted to synthetic tick density data to assess their ability to recover true parameter values. In each simulation, a set of parameter values was randomly selected from the respective prior distributions, and nymph and adult tick density data were synthesized based on this parameter set for dates corresponding to the observed tick density data. The model was then fitted to the synthetic data. These steps were repeated across 85 simulations to obtain the following statistics for each parameter.

First, the mean and ±1 standard deviation of the differences between the true parameter values (used to generate the synthetic data) and the posterior modes (obtained from model fitting) were calculated to assess biases in the posterior estimates. Second, the percentages of simulations with which the true parameter value fell within the 95% HDI of the posterior distribution was used to evaluate how well the posterior estimates captured the true parameter values.

While the assessment indicated that model T.12 can effectively recover parameter values used to generate synthetic tick density data, parameters related to early life stages, including oviposition ($\Theta_O$) and hatching ($\Theta_H$), for which density data were unavailable, exhibited a relatively wide range of bias from the true parameter values (Fig. S2C). When the daily oviposition rate was modelled using values inferred from controlled laboratory conditions (as in model T1.8 in Table 1), the bias range for $\Theta_H$ decreased significantly (Fig. S2C), suggesting potential influence of larva (or egg) density data on model fitting. However, while model T.12 allowed a greater level of uncertainty for these parameter values, it still suggests parameter estimates for all developmental processes based on field conditions and with a lowest DIC value. Therefore, it was chosen as the best-fitting model.

#### Generalised additive model (GAM) analysis and results

The GAMs models were fitted to the longitudinal questing tick density data, separately for the densities of questing nymphs and adults, using the *gam* function of the *mgcv* package(14). The best-fitting GAM was selected by comparing the fit of models that included random-effect terms for sampling site and sampling year, and combinations of cubic spline terms for the following climatic variables: day length, daily mean temperature, daily mean relative humidity, and daily mean saturation deficit. The selection of these climatic variables was conducted in a manual stepwise forward procedure.

For both questing nymph and adult densities, the best-fitting GAMs included cubic spline terms for day length, daily mean temperature, and daily mean relative humidity, in addition to random-effect terms for sampling site and sampling year (Table S2). For both nymphs and adults, day length showed the most profound effects on questing tick density, with increasing day length associated with higher questing density, whereas both daily mean temperature and relative humidity were negatively associated with questing density above certain levels (Figure S13).

The predictions for 2008–2042 were made based on the best-fitting GAMs and used as input data for the mechanistic model. Since some of those years were not included in the longitudinal questing tick density data, to which the best-fitting GAMs were fitted, the predictions were made without including the random-effect term for sampling year. $GAM\ fit_{l,s,d}$ represents the predicted tick density value for life stage $l$, sampling site $s$ on day $d$. These values were normalised, divided by the maximum predicted value observed across all sites during the study period, $GAM\ fit_{l_{max}}$. Finally, the normalised values were converted into daily questing rate values ($\tau_{l,s,d}$), multiplied by a scaling parameter δ.

Eq. 2 $\tau_{l,s,d} = \delta \frac{GAM\ fit_{l,s,d}}{GAM\ fit_{l_{max}}}$

δ was estimated as a single parameter, assuming that it was the same across the sites and life stages, given that the GAM fit values were derived after accounting for these differences. Therefore, it represented the maximum questing rate observed. Because GAM fit values were not available for larvae, it was assumed that larvae and nymphs exhibit identical questing rates, distinct from those of adults.

#### Mechanistic model parameters

For the association of the daily oviposition, hatching, and larva-to-nymph and nymph-to-adult moulting rates with climatic conditions assumed under Theme 1 null hypothesis, we used the inverse of the power-law curves estimated by Ogden, Lindsay (7). For Theme 1 null hypothesis, although Randolph, Green (15) also derived curves describing changes in the hatching and moulting rates of *Ixodes ricinus* at different temperatures based on laboratory observations, these curves were highly correlated with those derived by Ogden, Lindsay (7) and only available for hatching and moulting, but not oviposition rates. Therefore, we chose to use the Ogden, Lindsay (7) curves as the underlying functions for Theme 1 null hypothesis.

The daily background mortality rates, $m_0$, extracted from the literature were derived by through the observations of *I. ricinus* within containers placed in fields, for unfed and questing ticks in each life stage (3, 16). In the present study, these daily background mortality rates were assumed for unfed and questing ticks in each life stage (subscript $l$), linearly increasing with population size, becoming twice $m_0$ when population size reached its threshold value, $\omega_l$. It was assumed that the maximum questing nymph density observed across the study period was proportional to the threshold value for nymphs, allowing the estimation of the questing population size at this limit using a sampling fraction of 6.0% (17). Following this, the thresholds for larvae and adults were derived from the nymph threshold, taking into account theoretical survival probabilities between life stages (18). For unfed or questing ticks in life stage $l$ on site *s* and day *d*, with population of *N*, density-dependent mortality, $m_{1_{l,s,d}}$, was determined as follows:

Eq.4 $$m_{1_{l,s,d}} = m_{0_l}\left(1 + \frac{N_{l,s,d}}{\omega_l}\right)$$

For engorged ticks, mortality ($m_2$) was set to 1/365 to ensure that, on average, these ticks either progressed to subsequent life stages within one year after having their blood meal or died (overwintering no more than once).

#### Likelihood function for mechanistic model fitting

Briefly, after the tick population was simulated with a given set of parameter values, the predicted number of questing ticks per 100m$^2$ was calculated assuming a sampling fraction of 6.0% (17). Then, for site *s* and day *d*, the probability of the observed number of questing ticks per 100m$^2$ was assessed under a given probability distribution. Assuming an excess variability (overdispersion) in the observed data (19), we used a negative binomial distribution. The model was, therefore, fitted with the following likelihood function:

$$L = \prod_{s,d} NB(k_{N_q,s,d}; \mu_{N_q,s,d}, \emptyset_{N_q}) NB(k_{A_q,s,d}; \mu_{A_q,s,d}, \emptyset_{A_q})$$

For nymphs and adults, respectively, $k_{N_q,s,d}$ and $k_{A_q,s,d}$ were the observed and $\mu_{N_q,s,d}$ and $\mu_{A_q,s,d}$ were the predicted number of questing ticks per 100m$^2$ on site *s* and day *d*, and $\emptyset_{N_q}$ and $\emptyset_{A_q}$ were the overdispersion parameters for questing nymphs and adults, respectively. While the overdispersion parameters were assumed to be constant as a baseline, we also assessed whether allowing time-varying overdispersion would improve the fit of the best-fitting model by estimating separate overdispersion parameters for each astronomical or meteorological season.

For each model fitting process, the HMC sampling was iterated with multiple chains until convergence was considered achieved, based on visual inspection of the trace plots, Gelman-Rubin convergence diagnostic (<1.01), and the number of effective sample size (>500).
